# Nicotine in adolescence freezes dopamine circuits in an immature state

**DOI:** 10.1101/2023.10.28.564518

**Authors:** Lauren M. Reynolds, Aylin Gulmez, Sophie L. Fayad, Renan Costa Campos, Daiana Rigoni, Claire Nguyen, Tinaïg Le Borgne, Thomas Topilko, Domitille Rajot, Clara Franco, Fabio Marti, Nicolas Heck, Alexandre Mourot, Nicolas Renier, Jacques Barik, Philippe Faure

## Abstract

Nicotine use during adolescence is largely associated with negative long-term outcomes, including addiction to nicotine in adulthood. How nicotine acts on developing neurocircuitry in adolescence remains largely unknown, but may hold the key for informing more effective intervention efforts. We found transient nicotine exposure in early adolescence was sufficient for adult mice to show a marked vulnerability to nicotine. Brain-wide activity mapping showed that these mice had an enhanced response to an acute nicotine injection and widespread disruption of functional connectivity in comparison to controls, particularly within dopaminergic networks. Neurophysiological analysis further revealed that their ventral tegmental area (VTA) dopamine neurons show an immature basal plasticity signature and an adolescent-like imbalance in nicotine-induced activity between nucleus accumbens (NAc) and amygdala (AMG)-projecting pathways, known to respectively produce the reinforcing and anxiogenic effects of nicotine. The anxiogenic effect of nicotine is abolished in adult mice treated with nicotine in adolescence, strongly resembling the normal phenotype of young mice. Together these results suggest that nicotine exposure in adolescence somehow “froze” both their neural circuit and behavioral reaction to nicotine, carrying an adolescent-like vulnerability to the drug into adulthood. Finally, we are able to “thaw” the behavioral response to acute nicotine in adolescent-exposed mice by chemogenetically resetting the balance between the underlying NAc- and AMG-projecting dopamine circuits, restoring a mature anxiety-like response to acute nicotine. Together, our results highlight how diverse dopamine pathways can be impacted by experience in adolescence, and further suggest that the perseverance of a developmental imbalance between dopamine pathways may alter vulnerability profiles for later dopamine-dependent psychopathologies.

## Introduction

Smoking is a major contributor to disease burden worldwide, driven by addiction to nicotine^1^. The prognosis is particularly bleak for the up to 90% of adult smokers who began in adolescence^2^, as early onset drug use is associated with an elevated risk of addiction throughout the lifetime^3,4^. Indeed, epidemiological studies associate adolescent nicotine use with increased risk of not only longer and heavier smoking careers^5^, but also with the development of anxiety disorders and depression^6,7^, and with the problematic consumption of other drugs, such as alcohol^5^. Studies in animal models have focused to date largely on the differences in the rewarding effects of nicotine in adolescent vs adult rats and mice^8–14^, on how adolescent nicotine exposure can influence later measures of nicotine reward^15–23^, and on the long-term regulation of nicotinic acetylcholine receptors (nAChRs) and gene expression by nicotine in adolescence^24–27^. Despite significant interest in the effects of nicotine on the adolescent brain, little is known about how exactly nicotine affects the adolescent development of neural circuitry in order to increase vulnerability to later nicotine addiction and other psychopathologies.

Mature cognitive, emotional, and motivational behaviors emerge across adolescence in parallel to the development of their underlying neurocircuitry ^28–30^. Dopamine (DA) circuitry, in particular, is increasingly considered as a “plasticity system” where its structure and function is shaped by experience during development, creating adaptive behavioral profiles that can endure throughout the lifetime^31,32^. While this window may allow for performance optimization in the context of a specific environment, it also likely demarcates a period of increased vulnerability to environmental insult^33^. Nicotine acts directly on DA neurons through its action on nAChRs^34–36^, a family of pentameric ligand-gated ion channels. While the effects of nicotine on DA neurons are best studied in adult animals^37–39^, some evidence does suggest that both the immediate and enduring effects of nicotine on VTA DA neurons differ between adolescent and adult administration^40,41^.

However, these studies treat DA neurons as a single, homogenous population; whereas VTA DA neurons are increasingly recognized to belong to anatomically distinct circuits, to possess diverse molecular signatures, and to differ in their responses to external stimuli^42–48^. Indeed, nicotine simultaneously produces both reinforcing and anxiogenic behavioral effects^49–54^, which rely on distinct DA pathways: the activation of DA neurons projecting to the nucleus accumbens (NAc) produces reinforcement, whereas the inhibition of amygdala (AMG)-projecting DA neurons produces anxiety-like behavior^53^. Dopaminergic axons are still connecting to forebrain targets in adolescence^33^, and are sensitive to disruption by other psychostimulant drugs of abuse^55–57^; however whether and how nicotine may alter the development of distinct DA circuits in adolescence to drive later addiction vulnerability remains unknown.

## Results

### Exposure to nicotine during adolescence, but not during adulthood, enduringly increases vulnerability to nicotine use

To model nicotine exposure in the period when human adolescents are most likely to start smoking or using e-cigarettes^58,59^, we focused on comparing exposure to nicotine early in adolescence in mice (∼PND21 – PND 28)^33^ with exposure in adulthood (> PND60). To this end, male mice underwent a one-week exposure to nicotine (NIC, 100 µg/ml in 2% saccharin) or to 2% saccharin only (SAC) in their home cage drinking bottle either in early adolescence or in adulthood. Mice were always group housed for this pretreatment period, and for the following treatment washout period of five weeks, which allowed the adolescent-treated mice to reach adulthood. All mice were then tested for sucrose and nicotine consumption in a continuous-access free-choice oral self-administration paradigm (Fig 1A). Mice were weighed every two days when the bottle positions were changed, and their weight stayed stable over the course of the task (Sup Fig 1A). Mouse drinking behavior followed their diurnal behavioral profile, with the greatest drinking volumes shortly after the beginning of the dark cycle, regardless of the solution presented (Fig 1B). All mice showed a strong, dose-dependent preference for the sucrose-containing solution over water (Fig 1C), with similar overall volumes of liquid consumed (Sup Fig 1B), regardless of their pretreatment solution or age.

**Figure 1.**
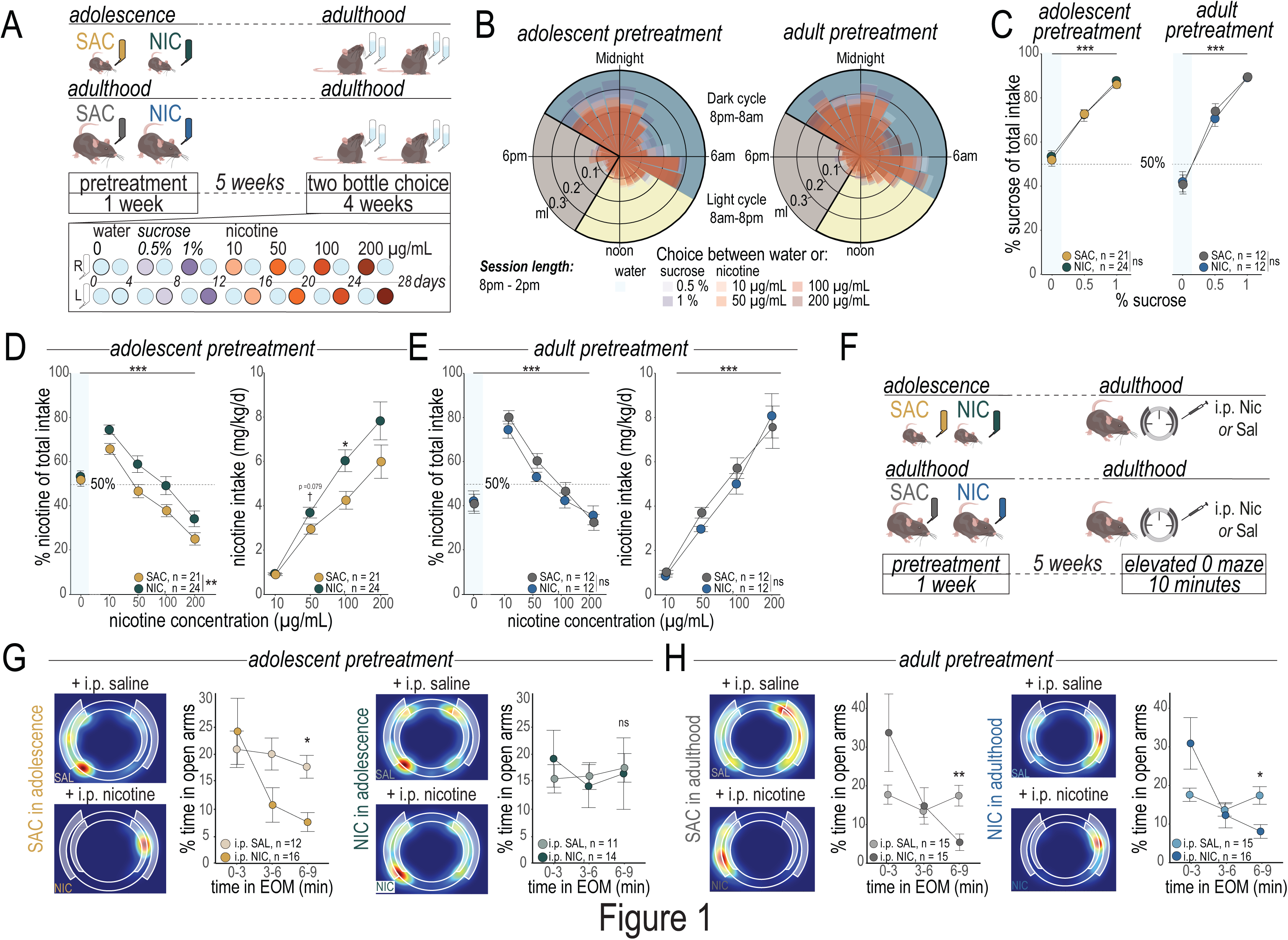
Enduring addiction-like vulnerability following brief exposure to nicotine in adolescence. (**A**) *Top:* Experimental timeline for mice treated with Nicotine (NIC; 100µg/mL nicotine in 2% saccharin) or Saccharin (SAC; 2% saccharin only) for one week in early adolescence (∼PND 21-PND 28) or in adulthood (starting at PND 75±15). All mice then underwent behavioral testing as adults, starting at PND 60 for the adolescent pretreated group and at PND 110 for the group pretreated as adults. *Bottom:* Timeline for the oral free-choice self-administration paradigm (Two Bottle Choice). (**B**) Actograms of drinking behavior over the duration of the Two Bottle Choice task for adult mice pre-treated in adolescence (*left*) or in adulthood (*right*). (**C**) Adult mice exposed to NIC in adolescence (*left*) or in adulthood (*right*) show equivalent preference for a sucrose solution as their SAC-treated counterparts. (**D**) Adult mice exposed to NIC in adolescence showed a higher percentage intake of the nicotine-containing solution over all treatment doses (*left*). Accordingly, the NIC-pretreated mice self-administered a higher daily dose of nicotine, with a significant difference at the 100µg/ml dose. (**E**) Mice pretreated with NIC in adulthood did not differ from SAC-treated controls in the percentage of nicotine solution consumed (*left*), nor by their daily dose of nicotine (*right*). (**F**) Experimental timeline. Adult mice treated with NIC or SAC in adolescence or in adulthood were tested for nicotine-induced anxiety-like behavior in the elevated 0 maze (EOM). (**G**) The anxiogenic properties of an acute nicotine injection are maintained in mice pretreated with SAC in adolescence (*left)* but blocked in adult mice that were treated with NIC in adolescence (*right)*. (**H**) Mice treated with NIC in adulthood show an equivalent anxiety-like response to acute nicotine (*right*) as their SAC-treated counterparts (*left*). All line graphs are presented as mean values ± SEM. ^†^ p < 0.08 *p < 0.05, **p < 0.01, ***p < 0.01, ns = not significant. Detailed Statistics are available in Table 1.

**Table 1.**
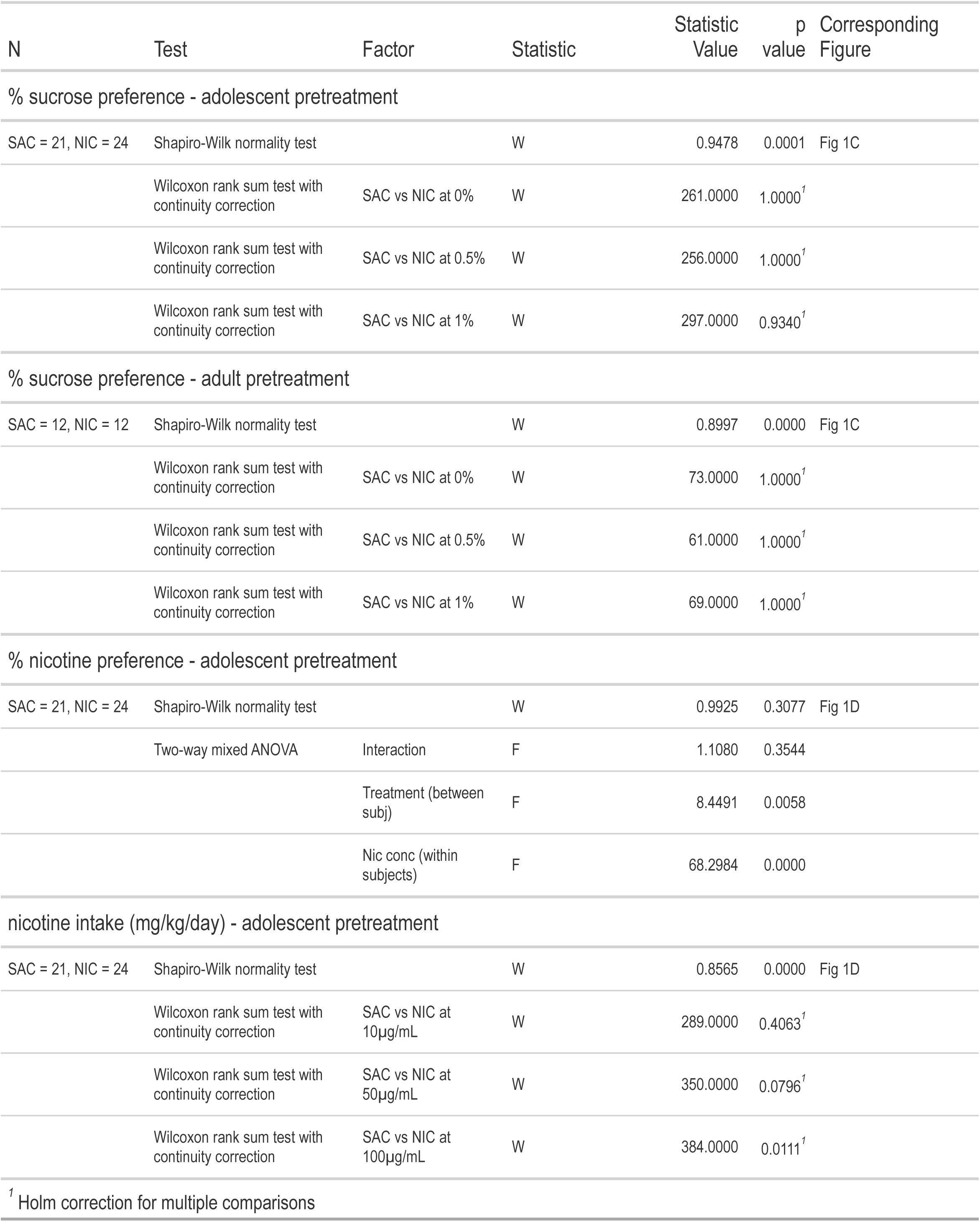
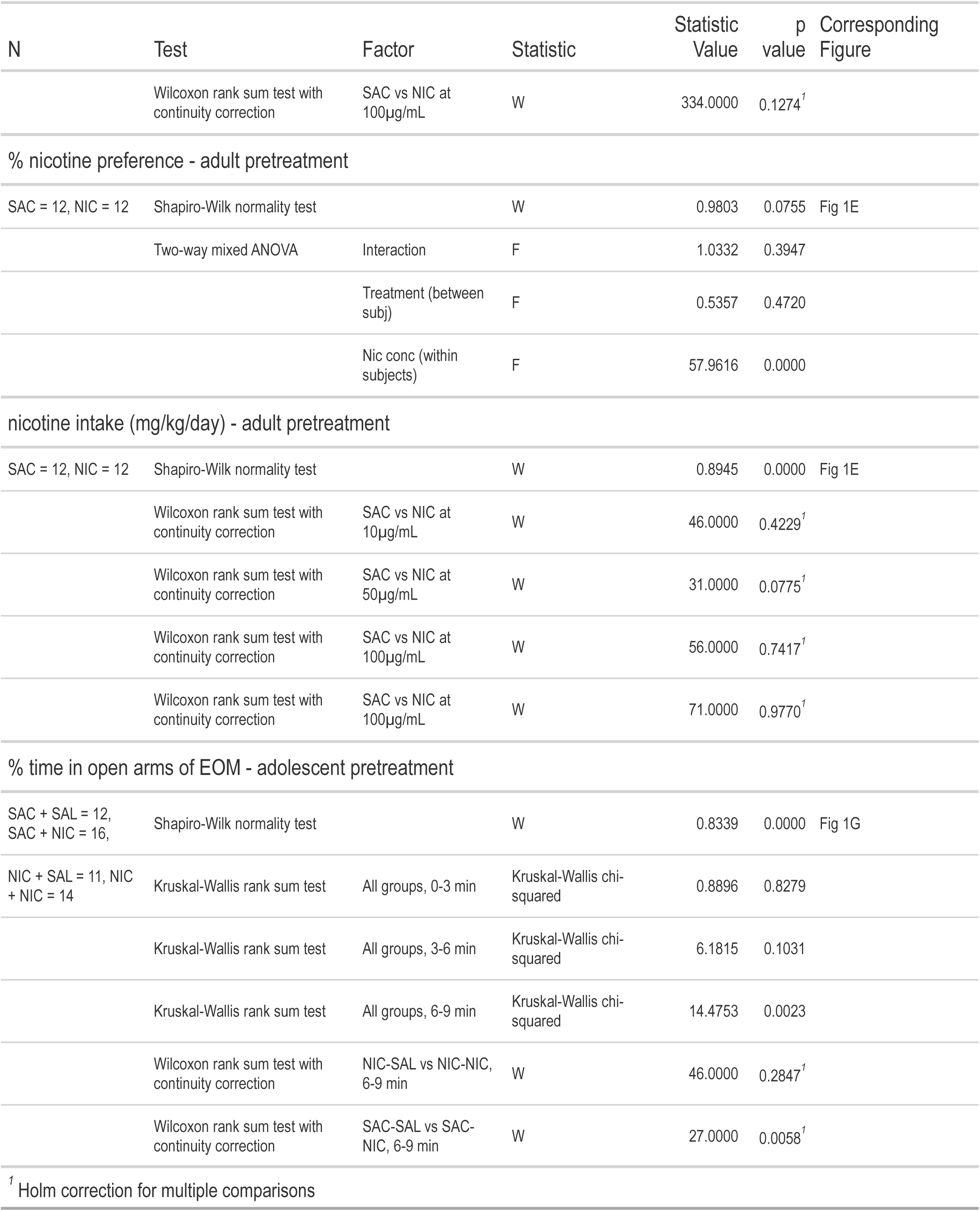
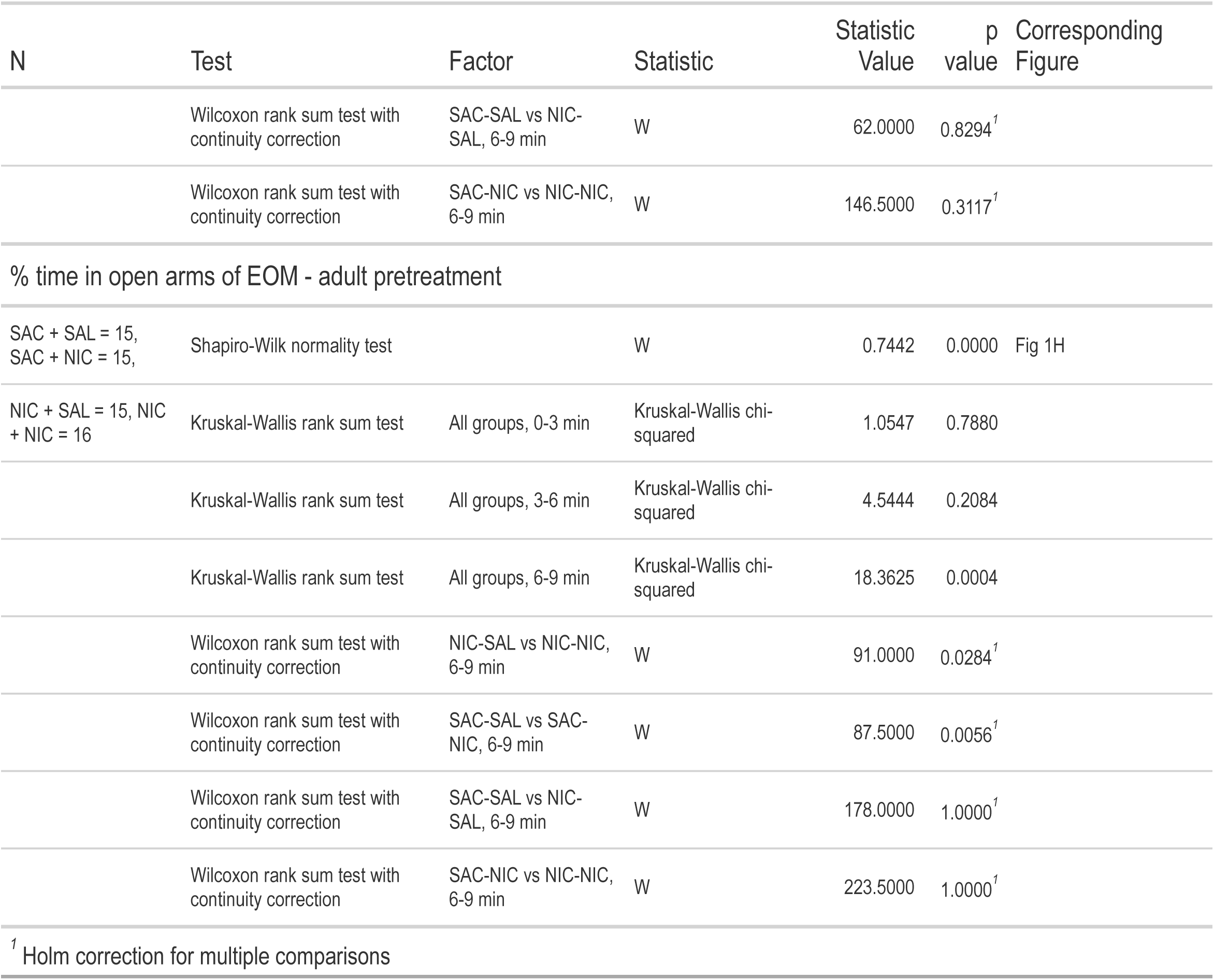
Detailed Statistics for Figure 1.

While rodents, and mice in particular, do not acquire intra-venous self-administration of nicotine as readily as with other stimulant drugs of abuse^60^, we found that mice voluntarily self-administered nicotine at all doses when tested in an oral nicotine intake paradigm (Fig 1D, 1E), with a strong effect of dose across all experiments indicative of a titration effect^61–65^. Mice showed the greatest preference for the nicotine solution at the 10µg/ml concentration, where it represented 65-80% of their liquid intake. Adult mice exposed to NIC in adolescence consumed more nicotine solution than their congeners exposed only to SAC in adolescence at all doses tested, indicative of vulnerability to nicotine use (Fig 1D). This did not result from side bias in consumption, as all mice were able to track the position of the nicotine or sucrose solution (Sup Fig 1C). In stark contrast, mice exposed to NIC in adulthood did not show different consumption behavior from their SAC-treated counterparts (Fig 1E), suggesting that this 1-week pretreatment regimen is too mild to produce enduring changes to the adult brain. Liquid intake across the nicotine sessions was stable across all groups of mice, and only the mice pretreated with NIC in adolescence showed a significant difference from their SAC-treated counterparts in the volume (mL) of nicotine solution consumed (Sup Fig 1D).

Nicotine has both reinforcing and anxiogenic properties^53^, both of which have been hypothesized to contribute to addiction liability. We thus next investigated how exposure to NIC in adolescence affects the anxiogenic properties of acute nicotine delivery in the elevated O-Maze (EOM, Fig 1F). While adult mice exposed to SAC in adolescence replicated the pronounced, time-dependent reduction of time spent in the open arms of the EOM seen after nicotine administration in naïve adult males^53^, exposure to NIC in adolescence abolished this anxiogenic effect (Fig 1G). However, the anxiogenic effect of acute nicotine injection was preserved in mice exposed to SAC or to NIC pretreatment as adults (Fig 1H). Adult mice exposed to NIC in adolescence, but not those exposed in adulthood, were also impervious to reductions in open arm entries (Sup Fig 1E) and suppression of locomotor activity (Sup Fig 1F) in response to acute nicotine injection.

Together, these results define an adolescent period where exposure to nicotine produces an enduring profile of altered response to later nicotine, featuring a reduction of its anxiogenic properties and an increase in voluntary consumption consistent with an “addiction-like” vulnerability. This finding further suggests vulnerability to nicotine is titrated along an age spectrum, where ongoing developmental processes create resilience to the drug as adolescents progress into adulthood.

### Nicotine in adolescence restructures nicotine-responsive networks in the adult brain

To dissect how this nicotine treatment in adolescence alters the functional organization of brain networks, we performed whole-brain activity mapping in a subset of mice that underwent testing for nicotine-induced anxiety-like behavior in the EOM (Fig 2A). Mice treated with NIC or SAC in adolescence or adulthood received an acute injection of nicotine or saline, were tested for anxiety-like behavior in the EOM, and were perfused ∼1 hour later at peak cFos expression. Their brains were then cleared using iDISCO, immunostained for cFos, and imaged on a lightsheet microscope^66^. Scans were automatically registered to the Allen Brain Atlas and cFos positive cells were quantified using the ClearMap analysis pipeline^67^. Adult mice exposed to NIC in adolescence had overall greater brain-wide activation in response to an acute injection of nicotine, with a significant increase over SAC treated counterparts in 129 regions (Fig 2B *left*). This included significant increases in cFos positive neurons in regions associated with addiction and addiction vulnerability, such as the nucleus accumbens (NAc), the prelimbic prefrontal cortex, and the pallidum; as well as in parts of the amygdala associated with anxiety and anxiety-like behavior (Anterior and Medial Amygdala). In contrast, mice that received NIC treatment as adults did not have major differences in cFos+ neuron expression from their SAC-treated counterparts (Fig 2B *right*); nor were major differences seen between any groups in response to saline injections (Sup Fig 2A, B).

**Figure 2.**
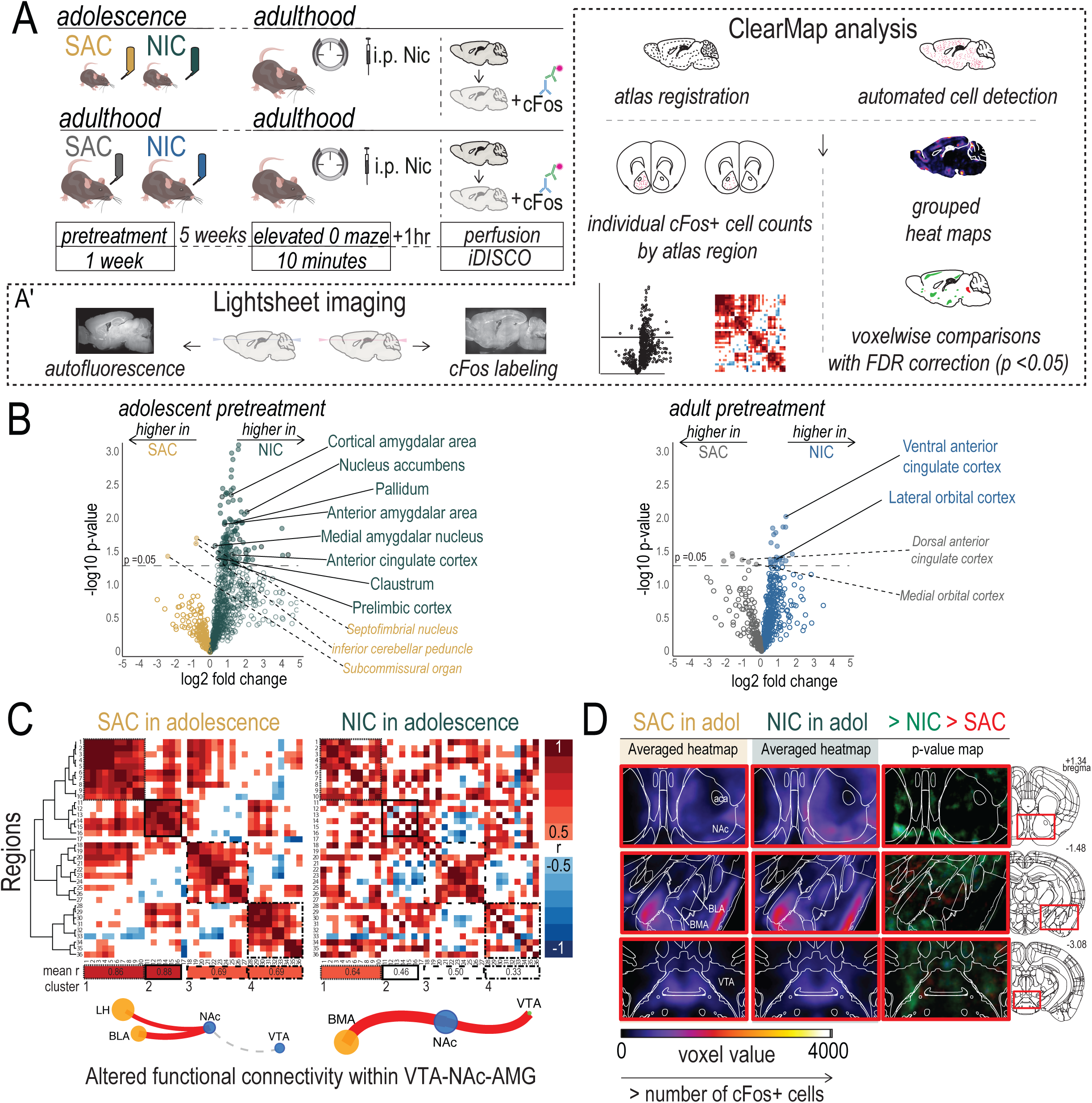
Nicotine in adolescence, but not in adulthood, reshapes brain-wide responsiveness to acute nicotine. (**A**) Experimental timeline for mice treated with Nicotine (NIC; 100ug/mL nicotine in 2% saccharin) or Saccharin (SAC; 2% saccharin only) in early adolescence or in adulthood. (**A’**) iDISCO brain clearing and Clearmap activity mapping pipeline. (**B**) (*Left*) Mice treated with NIC in adolescence show greater cFos activity across the brain, and in specific brain regions, following an acute nicotine injection when compared to their SAC-treated counterparts. Overall, 129 regions showed an increase in cFos activity in NIC-pretreated mice, defined as a fold change > 0 and a log p value > 1.3 (equivalent to p < 0.05). Notable regions of interest associated with anxiety response and/or response to nicotine have been highlighted. (*Right*) The activation profile of mice treated with NIC in adulthood is not substantially different from their SAC-treated counterparts. (**C**) Correlation matrices of relationships between cFos cell numbers in response to nicotine injection. Activation across brain regions in SAC-pretreated mice (*left*) is highly correlated, and hierarchical clustering organizes these regions into 4 distinct modules with the strongest inter-region relationships. In NIC-pretreated mice (*right*), correlations between regions are less strong. *Bottom*, community analysis on networks formed from these correlation matrices indicate significant reorganization of the VTA-NAc-AMG connections. (**D**) Voxel-by-voxel analysis of these regions reveals increases in nicotine reactivity within the VTA and DA terminal regions. Green or red voxels indicate a p value of < 0.05 after FDR correction.

To perform a more in-depth network analysis, we first identified a list of 36 brain regions implicated in anxiety behavior and nicotine response^68–72^. Correlation matrices show that nicotine-responsive regions are highly correlated in SAC-pretreated animals and can be organized into 4 modules with an average pearson correlation of 0.69 or more (Fig 2C *left*). When the same relationships between these regions are probed in NIC pretreated mice, a global disruption of correlated activity is apparent (Fig 2C *right*), with the strongest correlation in the same four modules at 0.64. The formation of strong functional correlations between these regions was less striking in mice of any pretreament condition that received saline injections before entering the EOM (Sup Fig 2 C, D), and mice that received SAC or NIC as adults had similar functional connectivity between the regions studied (Sup Fig 2E).

We then made network graphs from these nicotine-response correlation matrices, where each region was used as a node and each correlation as an edge, and we used Louvain community detection to organize these graphs into communities (Sup Fig 2G). We found that while the overall network structure, including community members and inter-community edges, were similar between the two pretreatment groups, there was a significant reorganization of the functional relationships between the NAc, the ventral tegmental area (VTA), and the basomedial and basolateral regions of the amygdala (Fig 2C *bottom*). We then confirmed that these areas showed significant voxel-by-voxel changes in the number of cFos+ cells in grouped comparisons, with more active cells seen in the NAc and VTA of NIC pretreated mice, and fewer active cells in the medial Amg of NIC pretreated mice in response to nicotine re-injection (Fig 2D). Mice that received SAC or NIC as adults did not show differences in the voxel-by-voxel comparison of grouped cell counts across these same regions (Sup Fig 2F). These results identify functional changes, particularly within dopaminoceptive networks, present in adult animals following exposure to NIC in adolescence, which may underlie the vulnerability to nicotine use we observed in these mice.

### Immature basal electrophysiological signature and exaggerated response to nicotine in dopamine neurons of adult mice exposed to nicotine in adolescence

While we discovered significant perturbation in DAergic functional networks following adolescent exposure to NIC in our brain-wide activity analysis, including changes in activation in the NAc and AMG regions, we did not find changes in DA bouton density in these terminal regions (Sup Fig 3A-D). Thus, we next assessed the electrophysiological profiles of DA neurons (Fig 3A) to elucidate whether network changes may result from altered DA neuron responsivity to nicotine. The peak amplitude of nicotine currents did not differ between mice exposed to NIC or SAC in adolescence (Fig 3B *left*) or adulthood (Fig 3B *right*), suggesting that nicotine signaling through nAChRs present on DA neurons remains intact following adolescent exposure and that altered response to nicotine in adults exposed during adolescence cannot be explained solely by differences in nAChR receptor composition or density on DA neurons. While there were no differences in presynaptic functioning of excitatory synapses (Sup Fig 3E), we found striking differences in glutamatergic input to the VTA (Sup Fig 3F) and post-synaptic plasticity at DA neurons only of adult mice exposed to nicotine in adolescence. These mice showed a reduction in AMPA/NMDA ratio in comparison to SAC-treated controls (Fig 3C *left*) and an increase in the weighted NMDA decay time constant (1w). These alterations to glutamatergic plasticity onto VTA dopamine neurons of adult mice more closely resemble conditions observed in young brains^40,42,73–75^, suggesting that nicotine exposure in adolescence is fixing plasticity mechanisms in an immature state.

**Figure 3.**
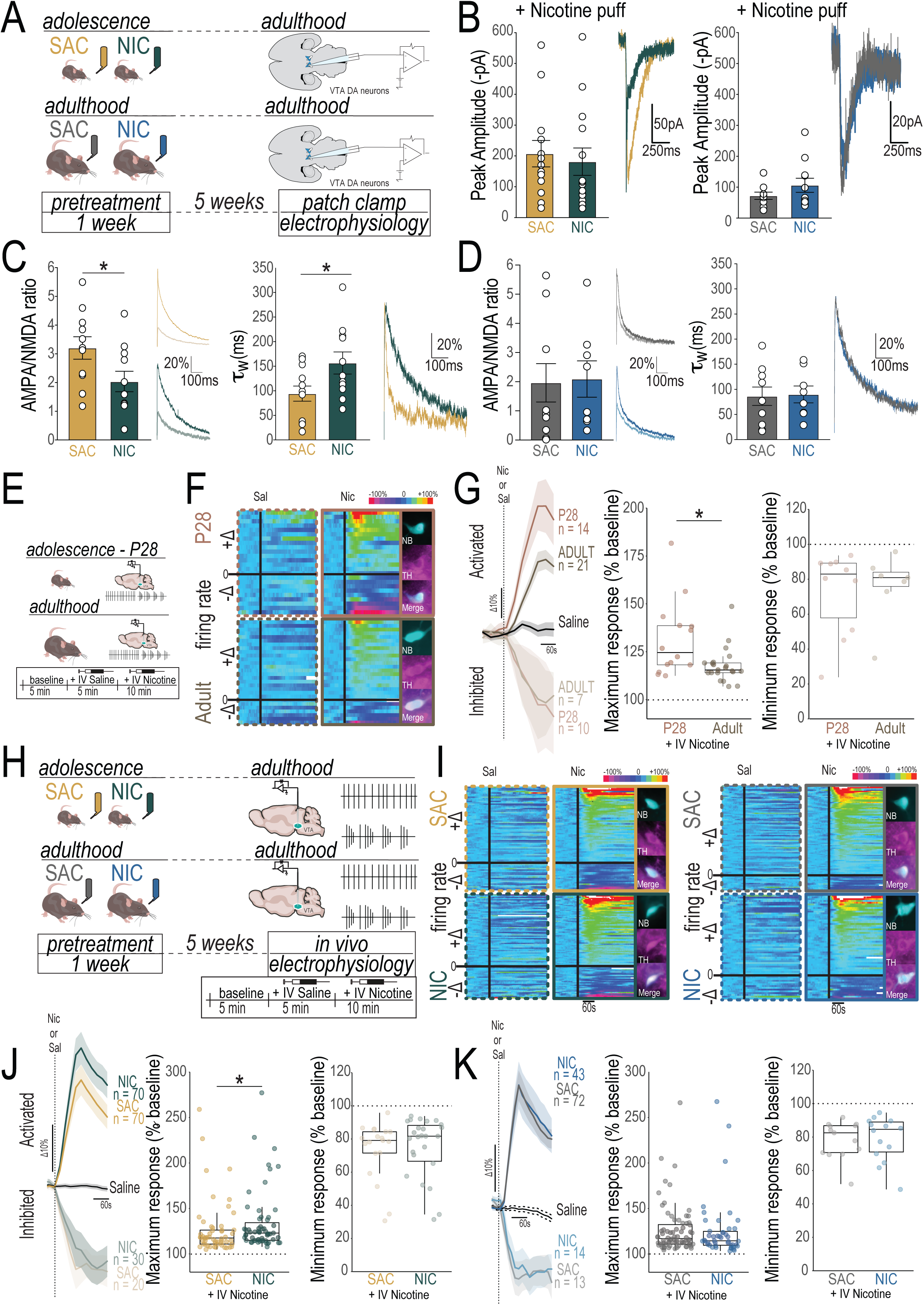
Persistent immature neurophysiological signature of VTA dopamine neurons in mice exposed to NIC in adolescence. (**A**) Experimental design for patch clamp experiments. (**B**) *Left*: No difference in peak amplitude of nicotine current between adult mice that received NIC or SAC as adolescents (NIC N = 2 mice, n = 14 neurons; SAC N = 2 mice, n = 13 neurons). *Right*: No difference in peak amplitude of nicotine current between adult mice that received NIC or SAC as adults (NIC N = 5 mice, n = 11 neurons; SAC N = 5 mice, n = 9 neurons). (**C**) *Left*: AMPA/NMDA ratio was decreased in mice that received NIC as adolescents (NIC N = 4 mice, n = 11 neurons; SAC N = 4 mice, n = 11 neurons). *Right*: NMDA current decay (weighted 1) was increased in mice that received NIC as adolescents (NIC N = 4 mice, n = 11 neurons; SAC N = 4 mice, n = 11 neurons). (**D**) *Left*: No differences in AMPA/NMDA ratio were observed when mice received NIC as adults (NIC N = 4 mice, n = 8 neurons; SAC N = 5 mice, n = 9 neurons). *Right*: NMDA current decay (weighted 1) was the same between mice that received NIC or SAC as adults (NIC N = 4 mice, n = 8 neurons; SAC N = 5 mice, n = 9 neurons). (**E**) Single unit juxtacellular recordings were made in adolescent (P28) or adult (>P60) mice to measure DA neuron response to nicotine. (**F**) Neuron responses to saline (Sal) or nicotine (Nic) injection represented as changes from baseline activity in adolescent (P28) mice (*Top*) and adult (>P60) mice (*Bottom*). *Insets*: Example neurons were labeled with neurobiotin (NB) after recording and processed for tyrosine hydroxylase (TH) immunohistochemistry to confirm their dopaminergic identity. (**G**) Dopamine neurons of adolescent mice showed an increase in nicotine evoked activation in comparison with adult animals (*center*), with no difference in inhibition (*right*). (**H**) Experimental design: mice were exposed to NIC or SAC in adolescence or adulthood and single unit juxtacellular recordings of VTA dopamine neurons were made five weeks later. Recordings included a 5-minute baseline, a 5-min recording following an IV saline injection, and a minimum of 10 minutes of recording following an IV nicotine injection of 30µg/kg. (**I**) Neuron responses to saline (Sal) or nicotine (Nic) injection represented as changes from baseline activity in adolescent-treated mice (*left*) and adult-treated mice (*right*). *Insets* : Example neurons were labeled with neurobiotin (NB) after recording and processed for tyrosine hydroxylase (TH) immunohistochemistry to confirm their dopaminergic identity. (**J**) Dopamine neuron responses to nicotine in mice pretreated with nicotine in adolescence resemble those of adolescent mice, they showed an increase in nicotine evoked activation in comparison with SAC-treated counterparts (*center*), with no difference in inhibition (*right*). (**K**) Dopamine neurons of mice treated with NIC or SAC as adults did not differ in their maximum change in firing rate in response to IV nicotine in activated neurons (*left*) nor in their minimum firing rate of inhibited neurons (*right*). All bar graphs are presented as mean values ± SEM. Box plots include a box extending from the 25th to 75th percentiles, with the median indicated by a line and with whiskers extending from the minima to the maxima. *p < 0.05, **p < 0.01, ***p < 0.01. Detailed Statistics are available in Table 2.

**Table 2.** Detailed Statistics for Figure 3.

| N | Test | Factor | Statistic | Statistic Value | p value | Corresponding Figure |
| --- | --- | --- | --- | --- | --- | --- |
| Nicotine current - adolescent pretreatment |  |  |  |  |  |  |
| NIC = 2 mice, 14 neurons; SAC = 2 mice, 13 neurons | Shapiro-Wilk normality test |  | W | 0.8200 | 0.0017 | Fig 3B |
|  | Wilcoxon rank sum test with continuity correction | SAC vs NIC | W | 108.0000 | 0.4233 |  |
| Nicotine current - adult pretreatment |  |  |  |  |  |  |
| NIC = 5 mice, 11 neurons; SAC = 5 mice, 9 neurons | Shapiro-Wilk normality test |  | W | 0.8200 | 0.0017 | Fig 3B |
|  | Wilcoxon rank sum test with continuity correction | SAC vs NIC | W | 42.0000 | 0.5949 |  |
| AMPA/NMDA Ratio - adolescent pretreatment |  |  |  |  |  |  |
| NIC = 4 mice, 11 neurons; SAC = 4 mice, 11 neurons | Shapiro-Wilk normality test |  | W | 0.9603 | 0.4963 | Fig 3C |
|  | Welch Two Sample t-test | SAC vs NIC | t | -2.2008 | 0.0398 |  |
| Weighted Tau - adolescent pretreatment |  |  |  |  |  |  |
| NIC = 4 mice, 8 neurons; SAC = 4 mice, 11 neurons | Shapiro-Wilk normality test |  | W | 0.9474 | 0.2808 | Fig 3C |
|  | Welch Two Sample t-test | SAC vs NIC | t | 2.2914 | 0.0343 |  |
| AMPA/NMDA Ratio - adult pretreatment |  |  |  |  |  |  |
| NIC = 4 mice, 8 neurons; SAC = 5 mice, 9 neurons | Shapiro-Wilk normality test |  | W | 0.8319 | 0.0057 | Fig 3D |
|  | Wilcoxon rank sum test with continuity correction | SAC vs NIC | W | 39.5000 | 0.7727 |  |
| Weighted Tau - adult pretreatment |  |  |  |  |  |  |
| NIC = 4 mice, 8 neurons; SAC = 5 mice, 9 neurons | Shapiro-Wilk normality test |  | W | 0.9380 | 0.2949 | Fig 3D |
|  | Welch Two Sample t-test | SAC vs NIC | t | 0.1470 | 0.8851 |  |
| maximum activation - naïve animals |  |  |  |  |  |  |
| Adolescent = 14, Adult = 21 | Shapiro-Wilk normality test |  | W | 0.7851 | 0.0000 | Fig 3G |
|  | Wilcoxon rank sum test with continuity correction | Adolescent vs. Adult | W | 215.0000 | 0.0230 |  |

**Table 2** Detailed Statistics for Figure 3
| N | Test | Factor | Statistic | Statistic Value | p value | Corresponding Figure |
| --- | --- | --- | --- | --- | --- | --- |
| maximum inhibition - naïve animals |  |  |  |  |  |  |
| Adolescent = 10, Adult = 7 | Shapiro-Wilk normality test |  | W | 0.8089 | 0.0027 | Fig 3G |
|  | Wilcoxon rank sum test with continuity correction | Adolescent vs. Adult | W | 37.0000 | 0.8836 |  |
| maximum activation - adolescent pretreatment |  |  |  |  |  |  |
| NIC = 70, SAC = 68 | Shapiro-Wilk normality test |  | W | 0.6534 | 0.0000 | Fig 3J |
|  | Wilcoxon rank sum test with continuity correction | NIC vs SAC | W | 1748.0000 | 0.0072 |  |
| maximum inhibition - adolescent pretreatment |  |  |  |  |  |  |
| NIC = 30, SAC = 21 | Shapiro-Wilk normality test |  | W | 0.7946 | 0.0000 | Fig 3J |
|  | Wilcoxon rank sum test with continuity correction | NIC vs SAC | W | 262.0000 | 0.3150 |  |
| maximum activation - adult pretreatment |  |  |  |  |  |  |
| NIC = 45, SAC = 72 | Shapiro-Wilk normality test |  | W | 0.6465 | 0.0000 | Fig 3K |
|  | Wilcoxon rank sum test with continuity correction | NIC vs SAC | W | 1810.0000 | 0.2884 |  |
| maximum inhibition - adult pretreatment |  |  |  |  |  |  |
| NIC = 14, SAC = 13 | Shapiro-Wilk normality test |  | W | 0.9032 | 0.0158 | Fig 3K |
|  | Wilcoxon rank sum test with continuity correction | NIC vs SAC | W | 84.0000 | 0.7524 |  |

To determine if mice also maintain an adolescent-like response to nicotine injection after adolescent exposure, we first defined adult and adolescent responses to nicotine using in vivo single unit recordings in anesthetized mice (Fig 3E). Dopamine neurons of naïve adult male mice show opposing responses to intravenous nicotine administration, with >90% of neurons projecting to the nucleus accumbens (NAc) activated in response to nicotine, and the >85% of neurons projecting to the amygdala (Amg) inhibited by nicotine in a previous study^53^. We first replicated this same distribution in adult mice, with DA neurons (identified by their electrophysiological characteristics and confirmed with neurobiotin labeling in a subset of neurons) showing both excitatory (putative VTA-NAc circuit) and inhibitory (putative VTA-Amg circuit) responses to i.v. nicotine injection (Fig 3F, bottom). Next, we found that this pattern held true in adolescent mice as well, with DA neurons either activated or inhibited by nicotine both present in the VTA (Fig 3F, top). Dopamine neurons from naïve adolescent mice showed a higher maximum activation in response to nicotine injection when compared to naïve adults (Fig 3G), while there was no difference in the level of DA neuron inhibition by nicotine between the ages. The fact that DA neuron responses to nicotine in adolescent animals are “imbalanced”, or are stronger in activation of the VTA-NAc pathway, may represent a physiological vulnerability signature, in line with reports of greater rewarding effects of nicotine in adolescent animals^8–11,13,14,76^.

We next assessed VTA dopamine neuron firing in adult mice exposed to SAC or NIC during adolescence or during adulthood (Fig 3H). We found that all groups of mice showed a spectrum of response to nicotine as observed in naïve animals, with responses again ranging from strong activation to strong inhibition by nicotine (Fig 3I). Interestingly, mice exposed to nicotine in adolescence showed a stronger, adolescent-like, activation of DA neurons in response nicotine than their SAC treated counterparts, with no change in the magnitude of inhibition (Fig 3J). This effect was not observed in mice exposed to nicotine as adults (Fig 3K), suggesting that nicotine in adolescence promotes the persistence of an endogenous adolescent state when the effects of nicotine are skewed in favor of its rewarding effect.

### Restoring adult-like balance in nicotine-evoked DA signaling unmasks mature behavioral response

Our electrophysiological results raise the intriguing hypothesis that exposure to nicotine in adolescence “freezes” dopamine circuitry in an imbalanced, immature state, which promotes vulnerability to nicotine use. We next sought to test whether adolescent-like DA functioning would explain the differences that we observed in the behavioral response to nicotine following preexposure in adolescence. We thus tested naïve adult and adolescent mice for nicotine-induced anxiety-like behavior in the EOM to establish whether there is a basal difference between adolescent and adult behavior. In accordance with previous results^53^, naïve adult mice showed a decrease in time spent in the open arms following an i.p. injection of nicotine, but not after an injection of saline, indicating an anxiety-like response to nicotine (Fig 4A *center*). However, when mice were tested at PND 28 this anxiety-like response to nicotine was absent, as the mice showed no difference in time spent in the open arms after nicotine or saline injection (Fig 4A *right*), in line with results from adolescent rats^77^. As this strongly resembled the behavioral effect that we observed in mice treated with NIC in adolescence (Fig 1G), this result suggests that not only does nicotine in adolescence freeze both neural and behavioral response nicotine in an immature state, but that imbalance between nicotine-induced activation and inhibition of VTA dopamine neurons is possibly masking the anxiogenic effect of the drug.

**Figure 4.**
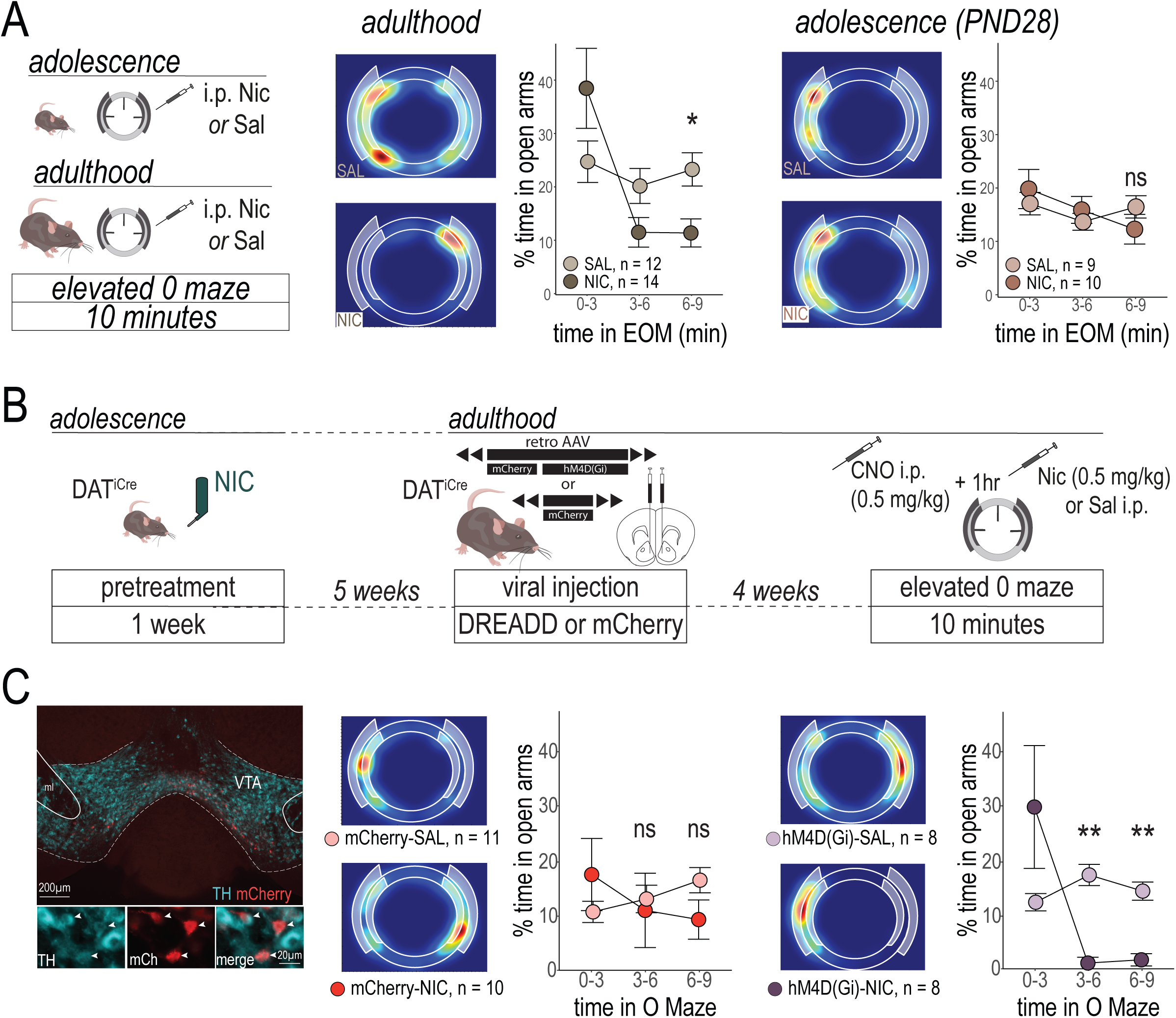
Chemogenetically dampening VTA-NAc circuit activity restores anxiety-like response to nicotine in adult mice exposed to nicotine in adolescence. (**A**) Experimental timeline for naïve mice (*left*). Adult mice spend less time in the open arms of the EOM following an injection of nicotine than an injection of saline (*center*). Adolescent mice show no difference in the time spent in the open arms of the EOM between a saline or nicotine injection (*right*). (**B**) Hierarchical clustering of all mice injected with nicotine before entering the EOM reveals that adult mice exposed to NIC in adolescence cluster only with the naïve adolescent mice (*more immature*), whereas mice that received NIC as adults cluster with the naïve adult mice and the mice exposed to SAC as adults (*more mature*). (**C**) Experimental timeline for DREADD intervention experiment. DAT^iCre^ mice were pretreated with NIC in adolescence, and then as adults they received an injection of a retroAAV hM4D(Gi) or control fluorescent-reporter virus into the NAc at the level of the medial shell. After 4 weeks, mice received an injection of CNO one hour before an injection of nicotine or saline and entering the EOM. (**D**) Retro AAV viruses were well expressed in VTA DA neurons of DAT^iCre^ mice following their behavioral testing (*left*). Mice that received a control virus replicated the effect of NIC in adolescence on WT mice, as mice never spent less time in the open arms of the EOM than their saline-treated counterparts (*center*). When VTA-NAc DA activity was reduced before nicotine injection, however, a mature behavioral response to nicotine injection, where mice spend less time in the open arms of the EOM, was restored (*right*). All line graphs are presented as mean values ± SEM. *p < 0.05, **p < 0.01, ***p < 0.01, ns = not significant. Detailed Statistics are available in Table 3.

**Table 3.** Detailed Statistics for Figure 4.

| N | Test | Factor | Statistic | Statistic Value | p value | Corresponding Figure |
| --- | --- | --- | --- | --- | --- | --- |
| % time in open arms of EOM - naïve adult animals |  |  |  |  |  |  |
| SAL = 12, NIC = 14, | Shapiro-Wilk normality test |  | W | 0.8878 | 0.0000 | Fig 4A |
|  | Wilcoxon rank sum test with continuity correction | NIC vs. SAC in 0-3 min | W | 104.0000 | 0.3159<sup>1</sup> |  |
|  | Wilcoxon rank sum test with continuity correction | NIC vs. SAC in 3-6 min | W | 49.0000 | 0.1519<sup>1</sup> |  |
|  | Wilcoxon rank sum test with continuity correction | NIC vs. SAC in 6-9 min | W | 35.0000 | 0.0377<sup>1</sup> |  |
| % time in open arms of EOM - naïve adolescent animals |  |  |  |  |  |  |
| SAL = 9, NIC = 10, | Shapiro-Wilk normality test |  | W | 0.9832 | 0.6108 | Fig 4A |
|  | Welch Two Sample t-test | NIC vs. SAC in 0-3 min | t | 0.6447 | 0.9308<sup>1</sup> |  |
|  | Welch Two Sample t-test | NIC vs. SAC in 3-6 min | t | 0.7492 | 0.9308<sup>1</sup> |  |
|  | Welch Two Sample t-test | NIC vs. SAC in 6-9 min | t | -1.2011 | 0.7408<sup>1</sup> |  |
| % time in open arms of EOM - DREADDS |  |  |  |  |  |  |
| MCH + SAL = 11,<br>MCH + NIC = 10, | Shapiro-Wilk normality test |  | W | 0.7376 | 0.0000 | Fig 4C |
| HM4 + SAL = 11,<br>HM4 + NIC = 14 | Kruskal-Wallis rank sum test | All groups, 0-3 min | Kruskal-Wallis chi-squared | 1.7264 | 0.6311 |  |
|  | Kruskal-Wallis rank sum test | All groups, 3-6 min | Kruskal-Wallis chi-squared | 18.0767 | 0.0004 |  |
|  | Kruskal-Wallis rank sum test | All groups, 6-9 min | Kruskal-Wallis chi-squared | 16.4805 | 0.0009 |  |
|  | Wilcoxon rank sum test with continuity correction | HM4-SAL vs HM4-NIC, 3-6 min | W | 1.0000 | 0.0033<sup>1</sup> |  |
|  | Wilcoxon rank sum test with continuity correction | MCH-SAL vs MCH-NIC, 3-6 min | W | 29.0000 | 0.1846<sup>1</sup> |  |
|  | Wilcoxon rank sum test with continuity correction | MCH-SAL vs HM4-SAL, 3-6 min | W | 56.0000 | 0.1846<sup>1</sup> |  |
|  | Wilcoxon rank sum test with continuity correction | MCH-NIC vs HM4-NIC, 3-6 min | W | 21.0000 | 0.3423<sup>1</sup> |  |

<sup>1</sup> Holm correction for multiple comparisons

Table 3 Detailed Statistics for Figure 4
| N | Test | Factor | Statistic | Statistic Value | p value | Corresponding Figure |
| --- | --- | --- | --- | --- | --- | --- |
|  | Wilcoxon rank sum test with continuity correction | HM4-SAL vs HM4-NIC, 6-9 min | W | 1.0000 | 0.0040<sup>1</sup> |  |
|  | Wilcoxon rank sum test with continuity correction | MCH-SAL vs MCH-NIC, 6-9 min | W | 28.0000 | 0.1368<sup>1</sup> |  |
|  | Wilcoxon rank sum test with continuity correction | MCH-SAL vs HM4-SAL, 6-9 min | W | 37.0000 | 0.1368<sup>1</sup> |  |
|  | Wilcoxon rank sum test with continuity correction | MCH-NIC vs HM4-NIC, 6-9 min | W | 18.0000 | 0.5915<sup>1</sup> |  |

<sup>1</sup> Holm correction for multiple comparisons

If an exaggerated, adolescent-like VTA-NAc DA neuron response to nicotine is masking the anxiogenic effects of the drug, dampening the activity of this circuit during nicotine administration could reveal the mature response. To test this possibility, we used a chemogenetic approach to exclusively decrease the excitability of VTA DA neurons of mice exposed to NIC in adolescence^78^, by expressing an inhibitory DREADD (hM4D(G_i_)) in a projection- and neurotransmitter-specific manner (Fig 4B). Mice that received the control virus recapitulated our previous results (Fig 1G) of an abolished anxiogenic response to nicotine, as mice that received nicotine in the EOM never spent less time in the open arms than their saline-treated counterparts (Fig 4C *center*). In contrast, dampening VTA-NAc DA activity with the DREADD virus caused the mice to spend dramatically less time in the open arms of the EOM after i.p. nicotine injection (Figure 4C *right*), an effect more closely resembling the response to nicotine in naïve adults (Fig 4A) and in adult mice treated with SAC in adolescence (Fig 1G). Both mCherry and DREADD injected mice that received CNO one hour before being tested with saline in the EOM showed similar levels of exploration of the open arms, in line with our other experiments, indicating that the CNO itself does not produce an anxiogenic or anxiolytic effect. The DREADD + nicotine mice were the only group tested that showed a significant anxiogenic response in the EOM, indicating that decreasing the excitability of VTA-NAc DA neurons restored the mature behavioral response to nicotine, effectively unmasking the anxiogenic effect of the drug. Together, this suggests that NIC in adolescence perpetuates a developmental signaling imbalance between discrete DA circuits in response to nicotine which masks the anxiogenic effect of the drug, leading to a vulnerability to increased nicotine taking in adult animals.

## Discussion

Nicotine use in adolescence remains a serious public health issue, with strong associations between early onset of use and later addiction^5^. Moreover, the widespread availability and acceptability of e-cigarette use has facilitated a startling increase in number of adolescent users and a concomitant decrease in their age of onset^58,79^. Understanding the mechanisms underlying this increased addiction risk can thus be useful for informing intervention efforts. We propose that exposure to nicotine in adolescence prolongs a developmental imbalance in dopaminergic circuitry, which, in turn, can promote vulnerability to nicotine use and addiction.

While the VTA has long been recognized as a heterogeneous structure, containing roughly 70-80% dopamine neurons, with the remaining 20-30% consisting of GABAergic and glutamatergic neurons^80–82^, studies on the function of dopaminergic neurons have only recently begun to address their diversity. Amongst themselves, dopamine neurons show significant physiological and molecular heterogeneity, as well as differences in their input-output wiring^42,83,84^. Increasingly, these diverse DA pathways are shown to play divergent roles in behavior, including responses to drugs of abuse^46,53,85,86^. Whether chronic drug exposure creates differential adaptations in distinct dopamine pathways to promote addiction remains an open question, with initial evidence indicating that chronic morphine can induce cell-body level morphological changes in neurons projecting to specific mesocorticolimbic regions^87^. We further suggest that enduring vulnerability to addiction following more subtle life experiences, such as early drug use or stress during development, may also result from pathway-specific adaptations in the DA system.

Our 1-week nicotine exposure paradigm is not sufficient to cause behavioral or physiological changes when this exposure occurs during adulthood. However, when this same exposure regimen is administered during adolescence, it produces enduring alterations to the response to a later nicotine challenge, priming the VTA-NAc dopamine pathway, specifically, to be more reactive to nicotine. This is in line with work showing that the acute and enduring effects of the adolescent brain to nicotine differ from those of adults. Notably, adolescent rodents are thought to be more sensitive to the rewarding effects of nicotine^8–14^, as well as less sensitive to its aversive effects, than adult animals^77,88,89^. The naturally occurring development of brain circuitry between adolescence and adulthood can thus be viewed as a protective mechanism against the deleterious effects of experience, and of nicotine use in particular. We found that the behavioral and physiological response to acute nicotine in adult mice pretreated with nicotine in adolescence strongly resembles the response of naïve adolescent mice. This finding suggests that nicotine exposure in adolescence blocks the protective normative maturation of nicotine-responsive systems and results in discordance between rewarding and aversive nicotine signals. We propose that this may be one mechanism by which nicotine use in adolescence creates an enduring vulnerability to later nicotine use and addiction. An imbalance between the rewarding and aversive effects of nicotine has indeed been posited as a mechanism driving the transition from casual use to addiction^61,90–92^.

Mesocorticolimbic DA circuits are known to undergo extraordinary growth and remodeling in the adolescent brain^33^, in contrast to other neuromodulatory systems which are largely established at the start of adolescence^93–95^. As such, the dopamine system is known to be sensitive to experience in adolescence, with significant effects of stress, social isolation, and drug use persisting until adulthood in many cases^32,33,96–98^. How exactly experience shapes dopaminergic function is still an open question, with possibilities including to advance, slow, or misroute normal maturational processes. Here, our evidence suggests that nicotine in adolescence “freezes” VTA-NAc and VTA-AMg DA circuits in their adolescent state. This stalling of developing DA circuits in response to experience is likely not limited to nicotine, or even drug exposure, as similar changes in glutamate plasticity onto VTA DA neurons have been reported after food insecurity in adolescence^99^. This raises important questions in the cross-sensitivity and additive relationship between stress and drug use in adolescence and adulthood. Other studies have reported pathological miswiring of DA circuits after psychostimulant exposure in adolescence^55,56,100^. While our evidence does not support the idea that nicotine in early adolescence leads to the global miswiring of dopamine axons, as the innervation patterns of dopamine axons in the NAc and AMG are unchanged, we find, rather, the persistent maintenance of adolescent-like responses to an acute nicotine challenge. Importantly, we also provide the first evidence that adult-like behavior can be restored in these developmentally “frozen” neural circuits by artificially re-calibrating neural responses to adult levels.

We focused our nicotine treatment on the early adolescent period (∼PND21-28) because not only does this correspond to the time of life when human adolescents are most likely to begin nicotine use^58,59^, but previous work in rodent models has also suggested that this is a particularly vulnerable time for DA development,^55,57^. Importantly, marked sex differences exist in adolescent developmental trajectories^101^, including dopamine development. Recent work has shown that adolescent periods of DA circuit vulnerability to experience differ between male and female mice, and that even when the immediate effects of an experience are the same, the enduring outcomes may differ dramatically due to sex- and/or age specific compensatory processes^55^, Indeed, nicotine receptor expression and function can be modulated by female sex hormones^102^, suggesting that even the immediate experience or effects of nicotine may be sex-dependent, adding to the complexity of addressing sex and developmental interactions in its effects. Ongoing work will address whether female mice show a similar or different pattern of adolescent DA circuit “freezing” by nicotine. Differential ages of vulnerability between sexes are also a key research arena, taking into account data from human studies showing that boys begin using nicotine younger than girls do^103^, and are more likely to escalate nicotine intake by transitioning from e-cigarette to cigarette use^104^.

Why adolescent nicotine users are more likely to continue to use nicotine, to have longer and heavier smoking careers, and to develop cross-drug addictions and/or psychiatric symptoms has largely remained unexplained at a mechanistic level. Here, we show that nicotine in adolescence locks the function of DA neurons in a persistent adolescent-like state, leading to exaggerated response to the rewarding effects of nicotine and a blunting of its negative, anxiogenic effects. This adolescent-like bias in behavioral responding may therefore facilitate the transition from casual use to addiction across the lifetime. Because we were able to “thaw” this response to nicotine - or restore adult-like functioning - with a chemogenic approach, our findings suggest that these effects of nicotine in adolescence may be reversible. Developing interventions that target restoring an appropriate balance between the positive and negative effects of nicotine use may have therapeutic applications.

## Acknowledgments

This work was supported by the Centre National de la Recherche Scientifique CNRS UMR 8246, INSERM U1130, the Foundation for Medical Research (FRM, Equipe FRM DEQ2013326488 to PF), the French National Cancer Institute (Grant TABAC-16-022, TABAC-19-020 and SPA-21-002 to PF), and French state funds managed by the ANR (ANR-20 NICADO to PF and JB, ANR-19-CE16-0028 Bavar to PF and NR). LMR was supported by a NIDA–Inserm Postdoctoral Drug Abuse Research Fellowship. CN was supported by a fourth-year PhD fellowship from the Biopsy Labex. We are grateful for support from the animal facilities (IBPS) and Otilia de Oliveira, Noemie Karakaplan-Dherbe and Emilie Tubeuf at ESPCI animal facilities.

## Author contributions

LMR and PF designed the study. LMR, AG, and CF performed the behavioral experiments. LMR, TT, and DRajot performed iDISCO and ClearMap experiments with support from NR. RCC and DRigoni performed ex vivo electrophysiological recordings with support from JB. LMR, SLF, CN, TLB, and FM performed in vivo electrophysiological recordings. LMR performed the surgeries and virus injections. LMR and AG performed quantitative neuroanatomy experiments with support from NH. LMR, AG, RCC, DRigoni, TT, JB, and PF analyzed the data. LMR wrote the paper with inputs from AM, FM, JB, and PF. All authors read and edited the manuscript. JB and PF secured the funding.

## Declaration of interests

The authors declare no competing financial interests.

**Supplementary Figure 1.**
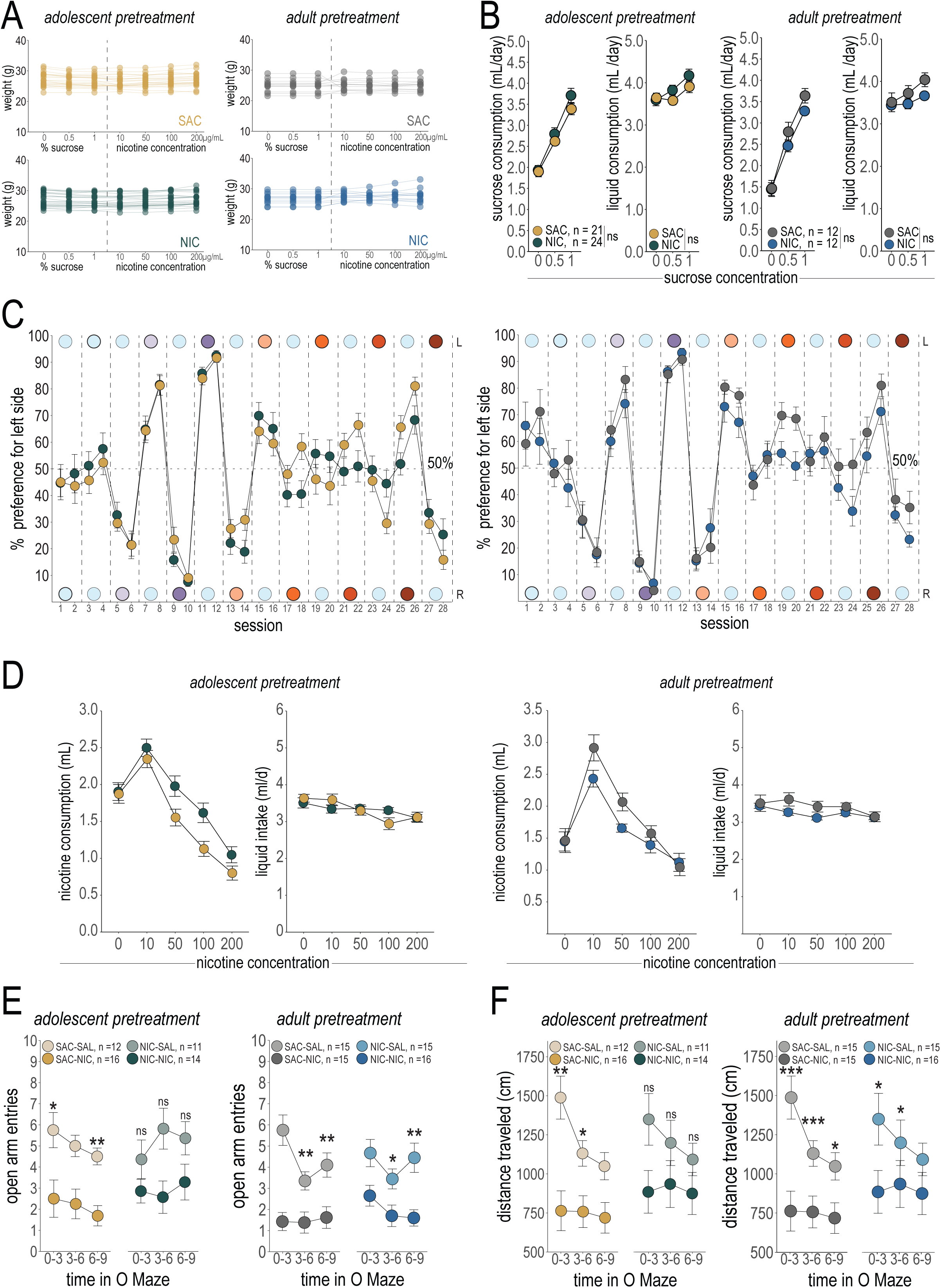
Supplementary data for Figure 1. (**A**) Weight measurements across the entire oral self-administration task for adolescent-pretreated (*Left*) and adult-pretreated mice (*Right*). (**B**) Sucrose solution and overall liquid consumption per day for adolescent-pretreated (*Left*) and adult-pretreated mice (*Right*) during the Sucrose nicotine oral self-administration testing. (**C**) Percent preference for left side bottle across the entire oral self-administration task for adolescent-pretreated (*Left*) and adult-pretreated mice (*Right*). (**D**) Nicotine solution and overall liquid consumption per day for adolescent-pretreated (*Left*) and adult-pretreated mice (*Right*) during the nicotine oral self-administration testing. (**E**) Open arm entries in the EOM task for adolescent-pretreated (*Left*) and adult-pretreated mice (*Right*). (**F**) Distance traveled (cm) in the EOM for adolescent-pretreated (*Left*) and adult-pretreated mice (*Right*). All line graphs are presented as mean values ± SEM. *p < 0.05, **p < 0.01, ***p < 0.01, ns = not significant. Detailed Statistics are available in Table S1.

**Supplementary Figure 2.**
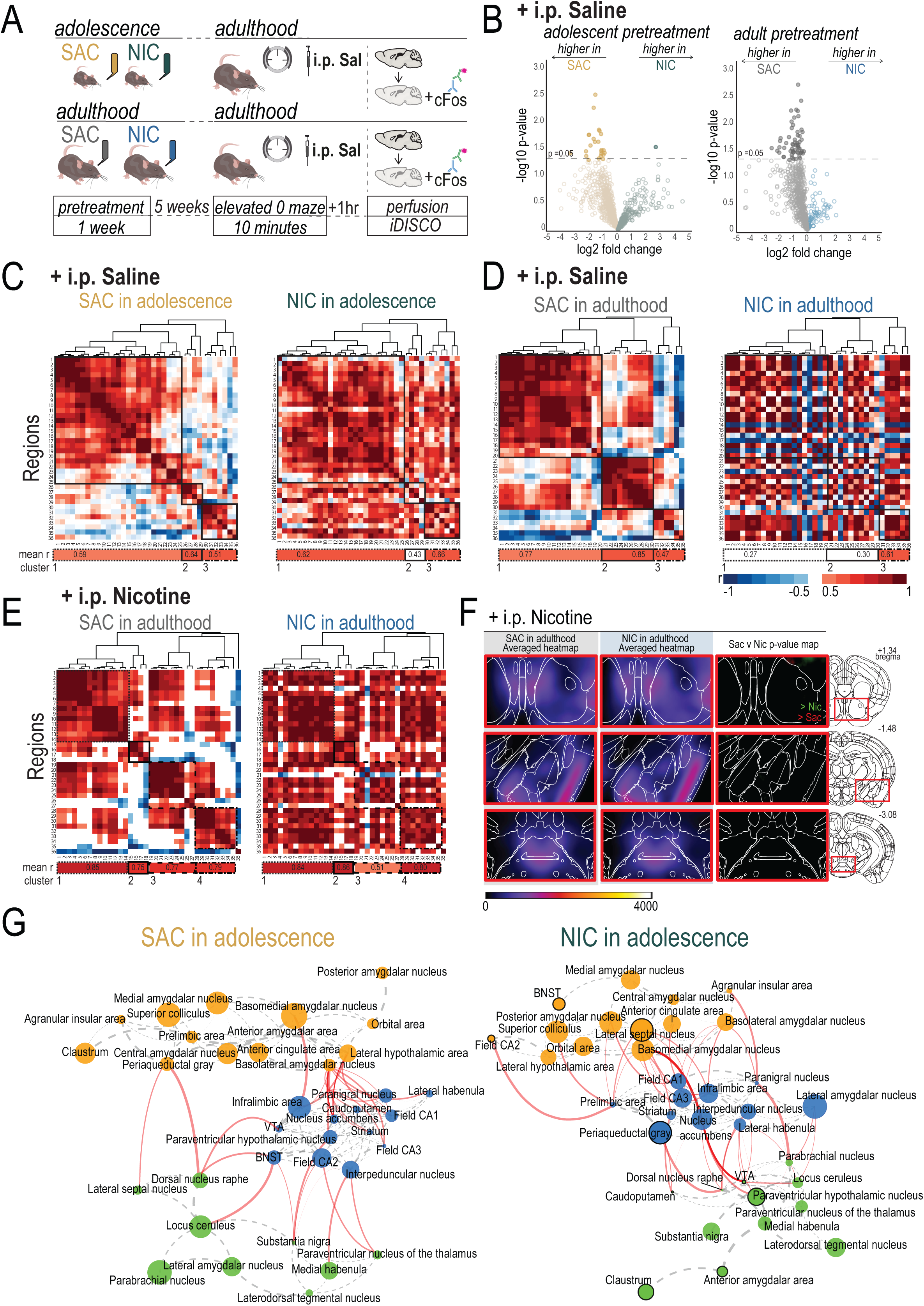
Supplementary data for Figure 2. (**A**) Experimental strategy. (**B**) Volcano plots showing differential cFos expression across brain regions following an injection of saline for adolescent-pretreated (*Left*) and adult-pretreated mice (*Right*). (**C-D**) Correlation matrices for relationships between cFos expression in brain regions following an injection of saline in adolescent-pretreated (**C**) and adult-pretreated mice (**D**). (**E**) Correlation matrices for relationships between cFos expression in brain regions following an injection of nicotine in adult-pretreated mice. (**F**) Voxel-by-voxel analysis reveals no change in reactivity to nicotine within the VTA and DA terminal regions in adult-pretreated mice. (**G**) Full networks derived from correlation matrices in Fig 2C and organized into communities with Louvain community analysis.

**Supplementary Figure 3.**
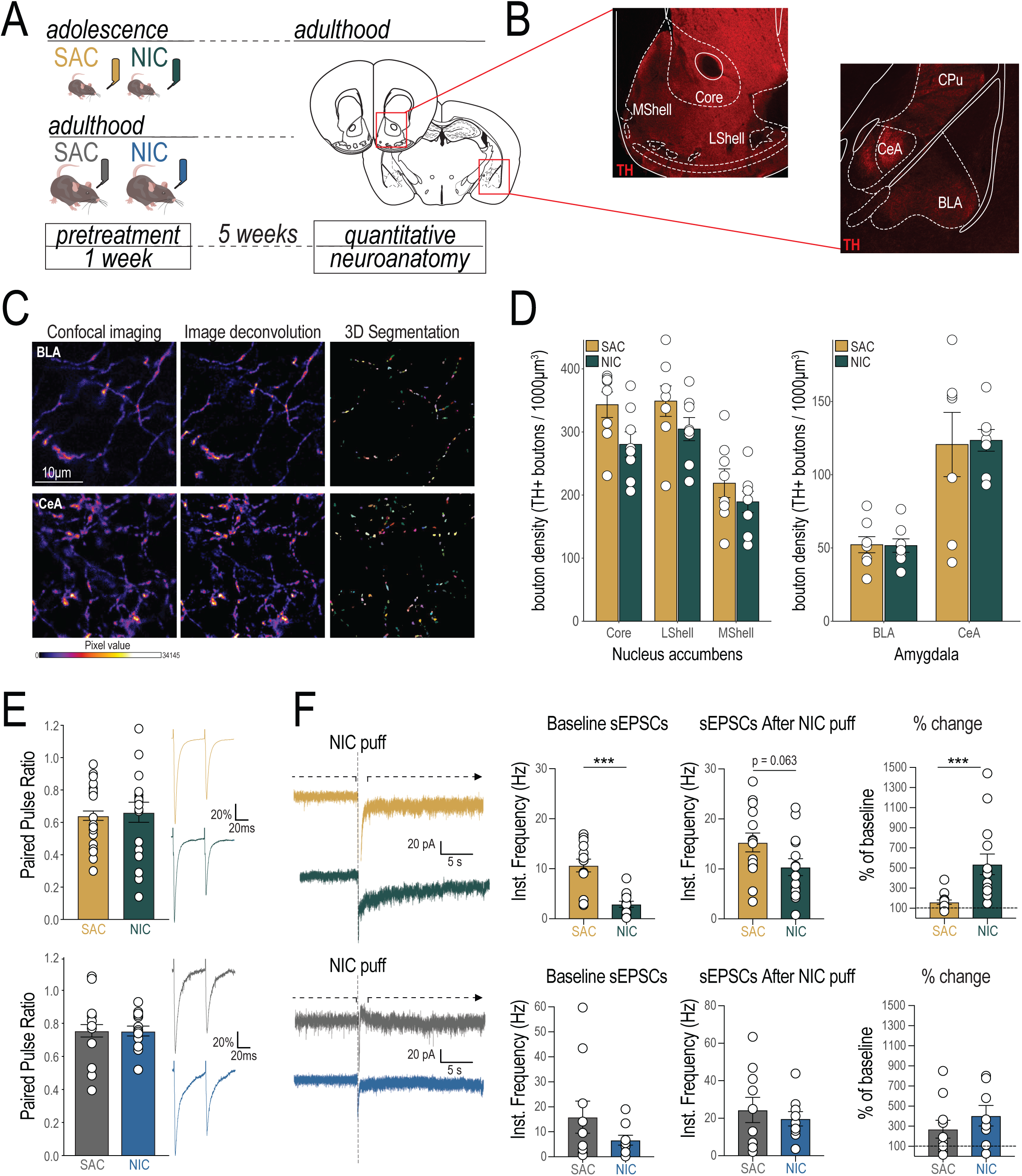
Supplementary data for Figure 3. (**A**) Experimental design for DA terminal density analysis. (**B**) Tyrosine Hydroxylase (TH)+ labeling in DA terminal regions. (Core = NAc Core, MShell = NAc Medial Shell, LShell = NAc Lateral Shell, CeA = Central Amygdala, BLA = Basolateral Amygdala, CPu = Caudate-Putamen). (**C**) Confocal image processing for bouton segmentation. (**D**) Bouton density counts across the NAc (*Left*) and the AMG (*Right*). (**E**) Paired pulse ratio measurements for VTA DA neurons of adolescent-pretreated (*Top*) and adult-pretreated mice (*Bottom*). (**F**) sEPSCs onto DA neurons measured before and after nicotine puff (*Left*) sEPSC frequency was reduced at baseline and after nicotine puff in adult mice pretreated with NIC in adolescence in comparison to mice pretreated with SAC (*Top row*), however NIC pretreated mice showed a greater augmentation (% of baseline) in response to the nicotine puff (*Right*). Mice pretreated with NIC in adulthood showed no differences from mice pretreated with SAC in adulthood (*Bottom row*). All bar graphs are presented as mean values ± SEM. *p < 0.05, **p < 0.01, ***p < 0.01. Detailed Statistics are available in Table S2.

**Table S1.**
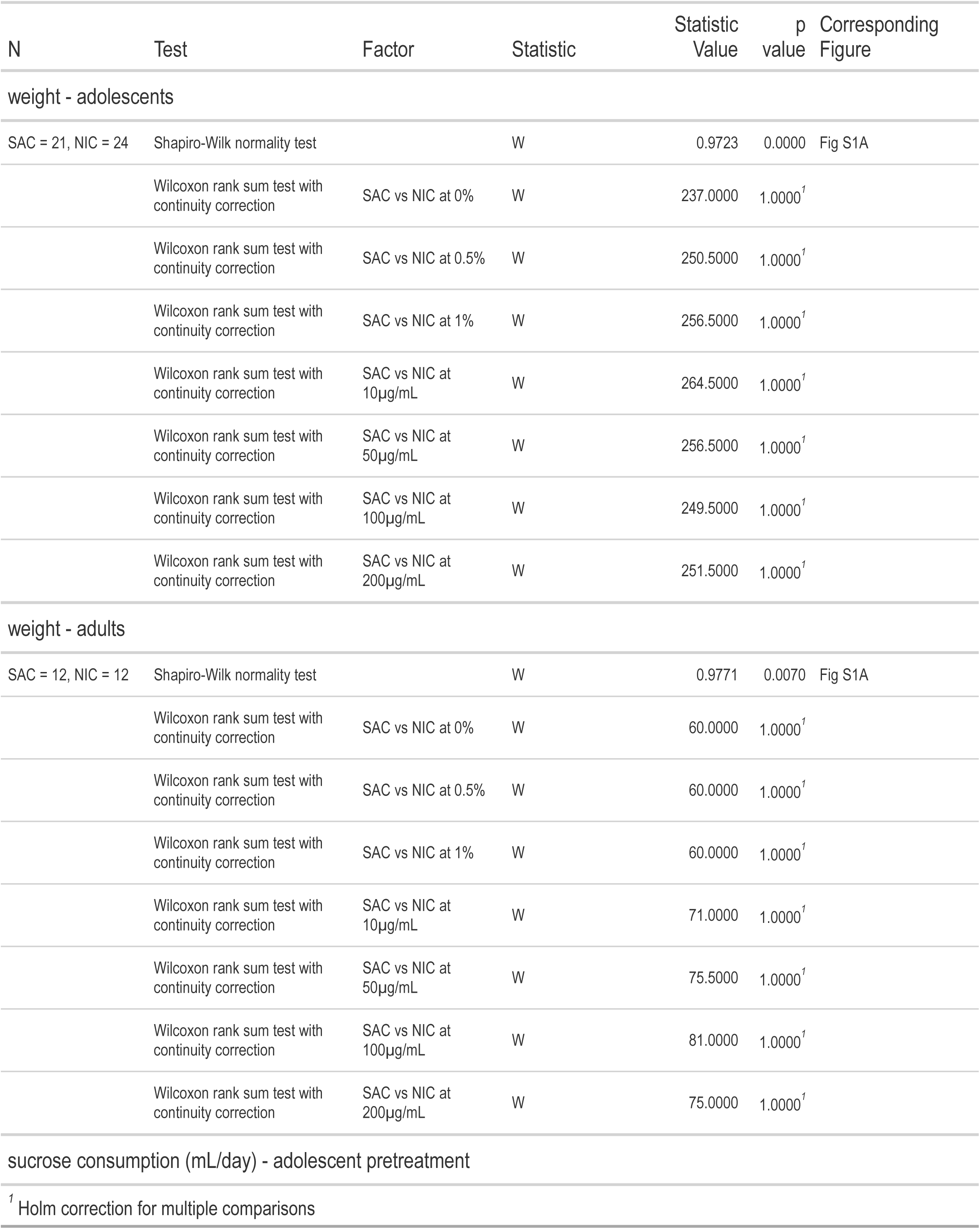
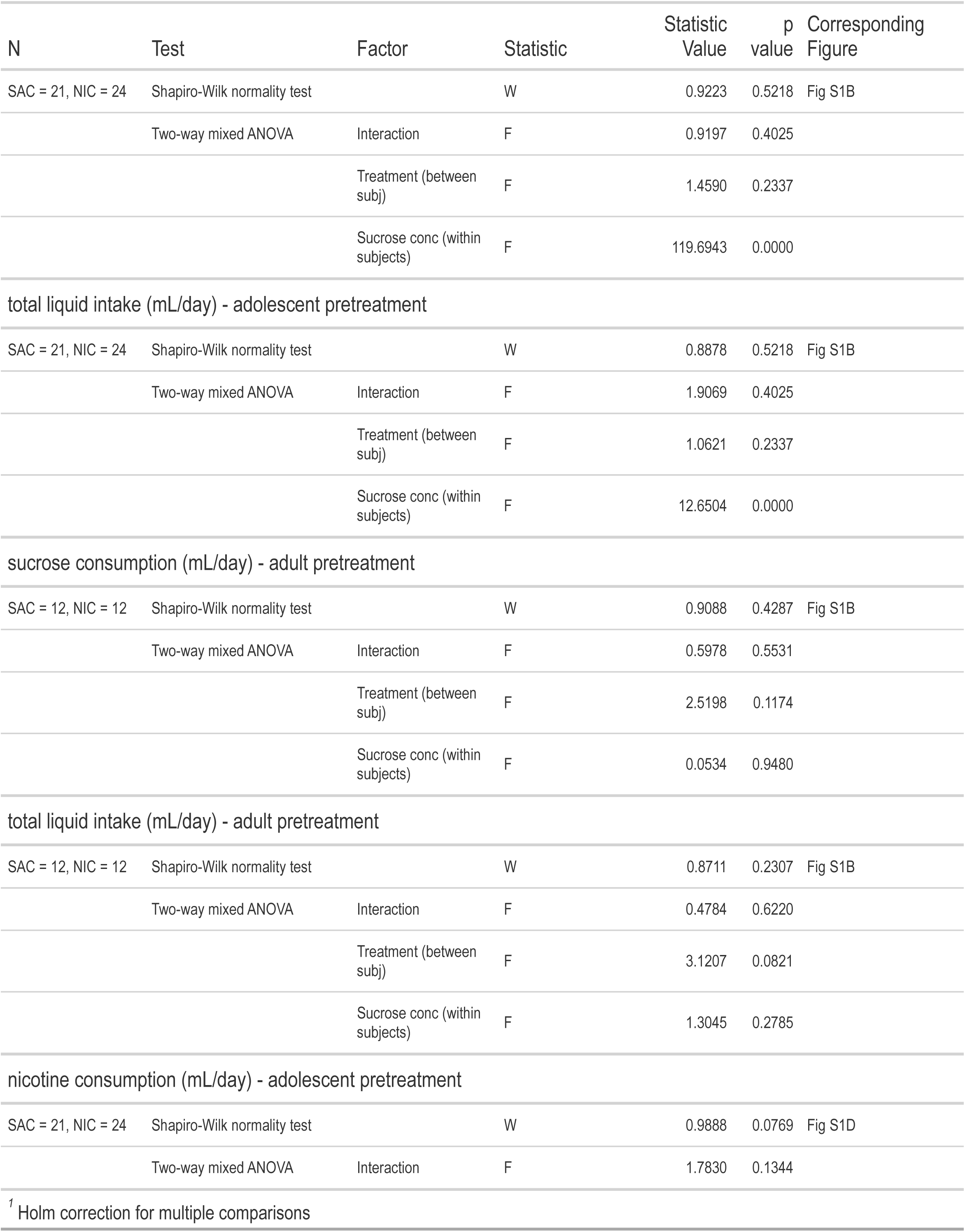
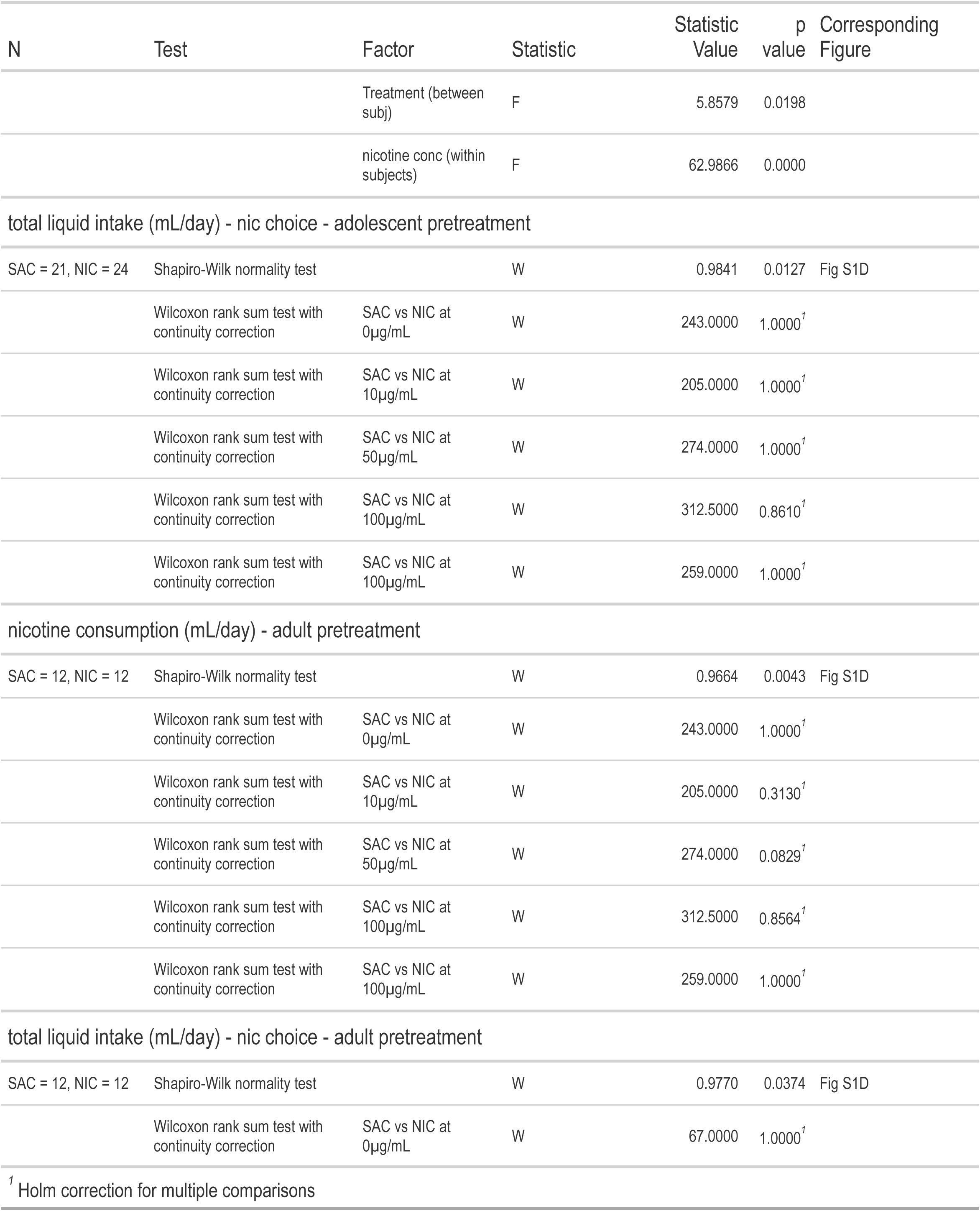
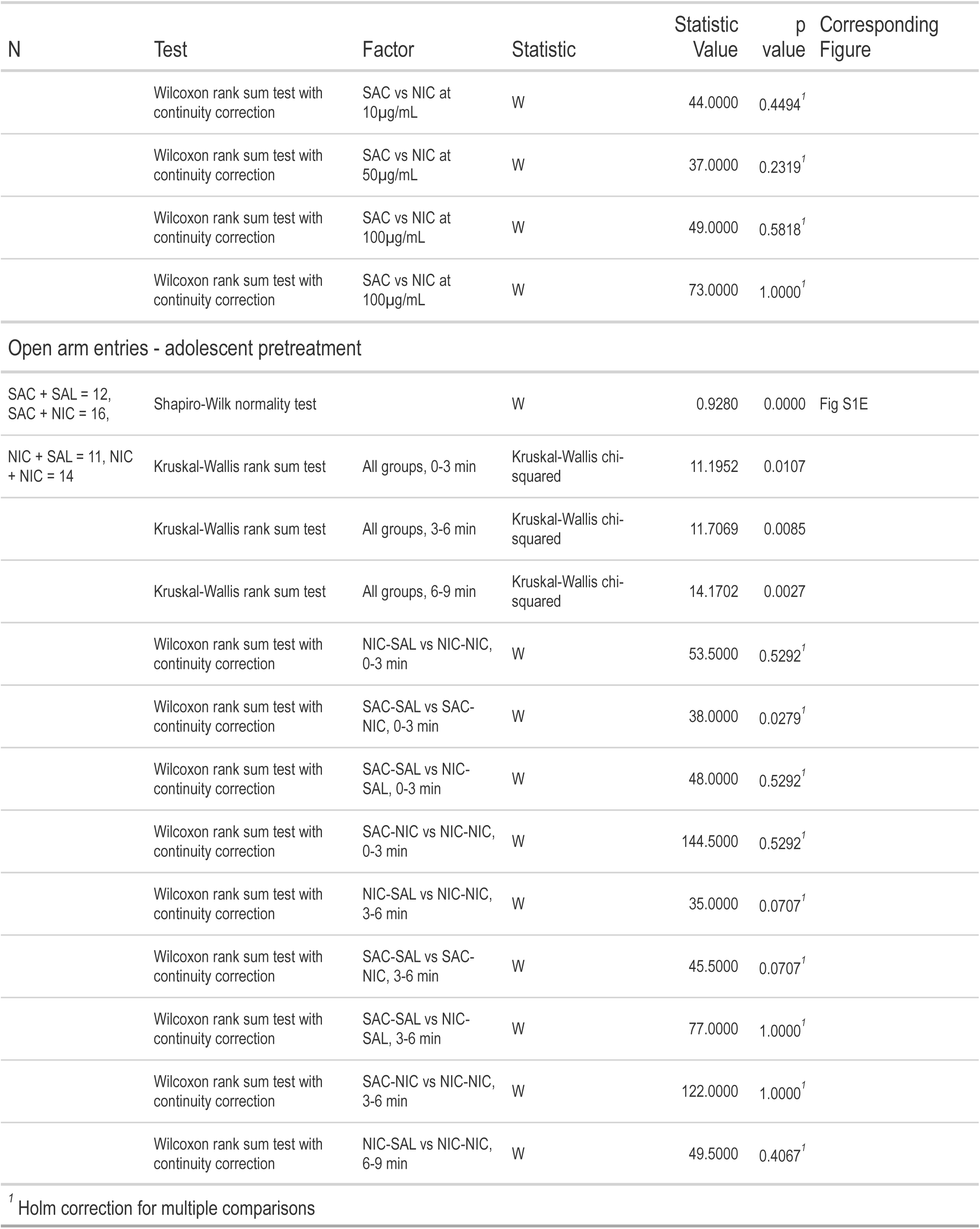
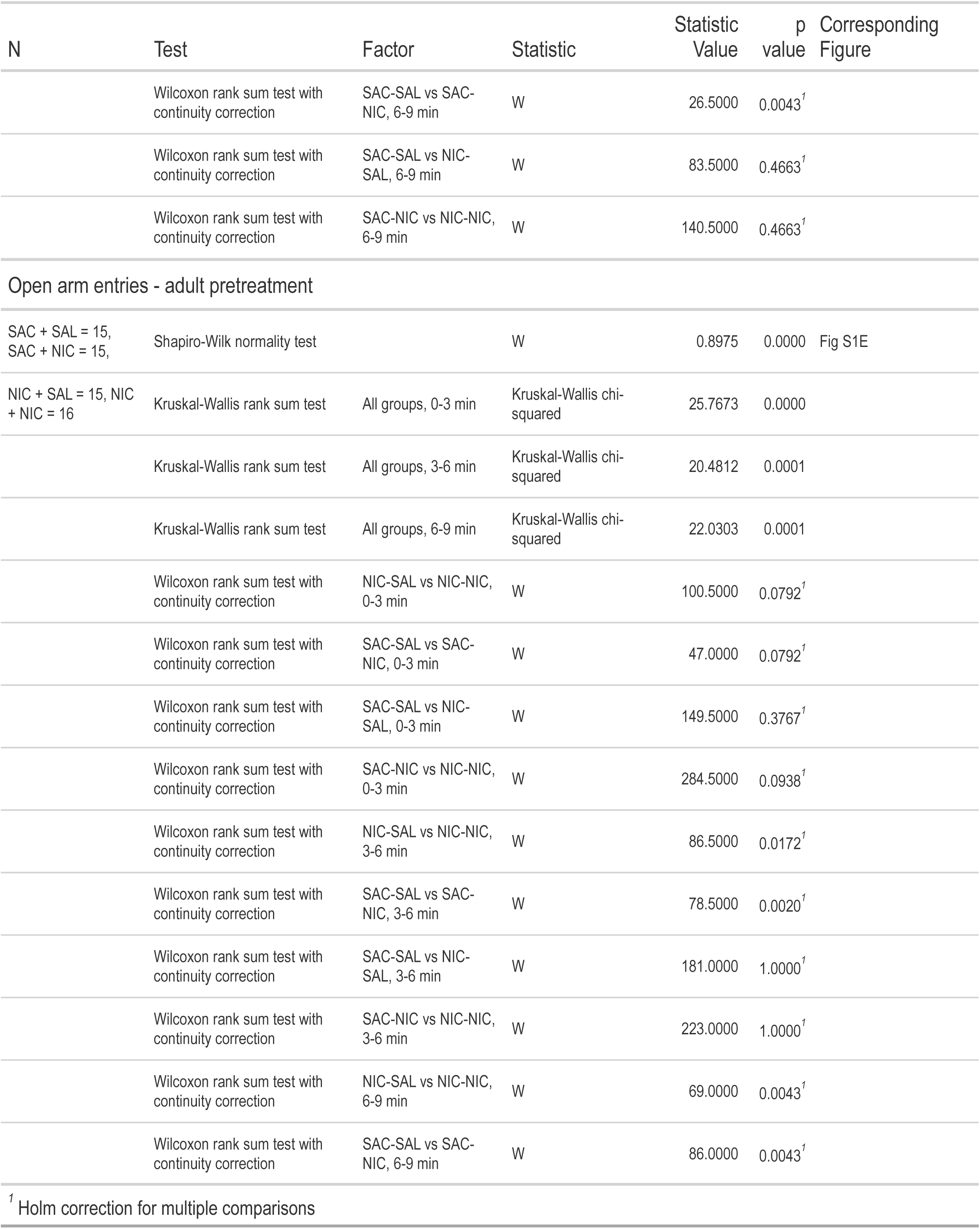
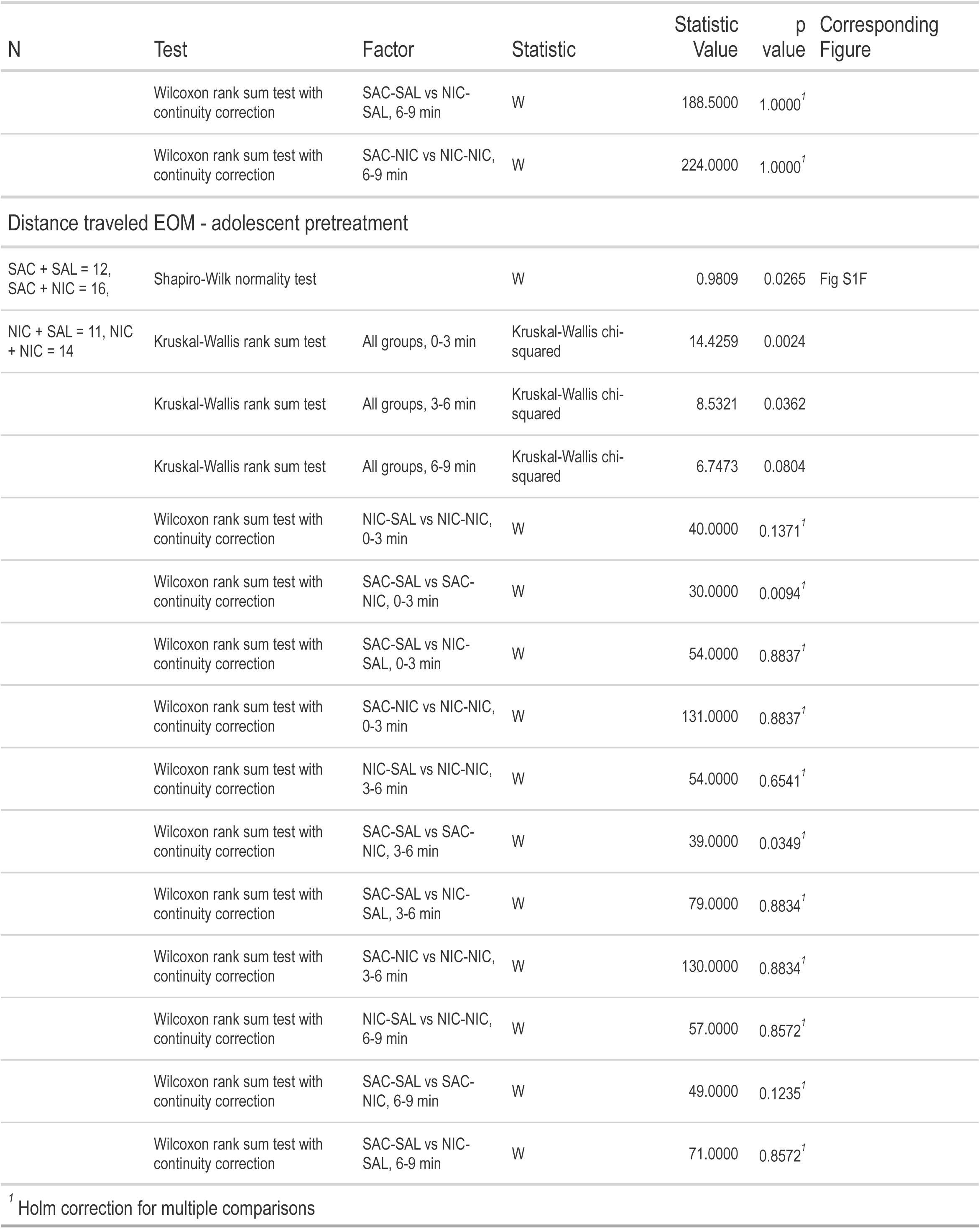
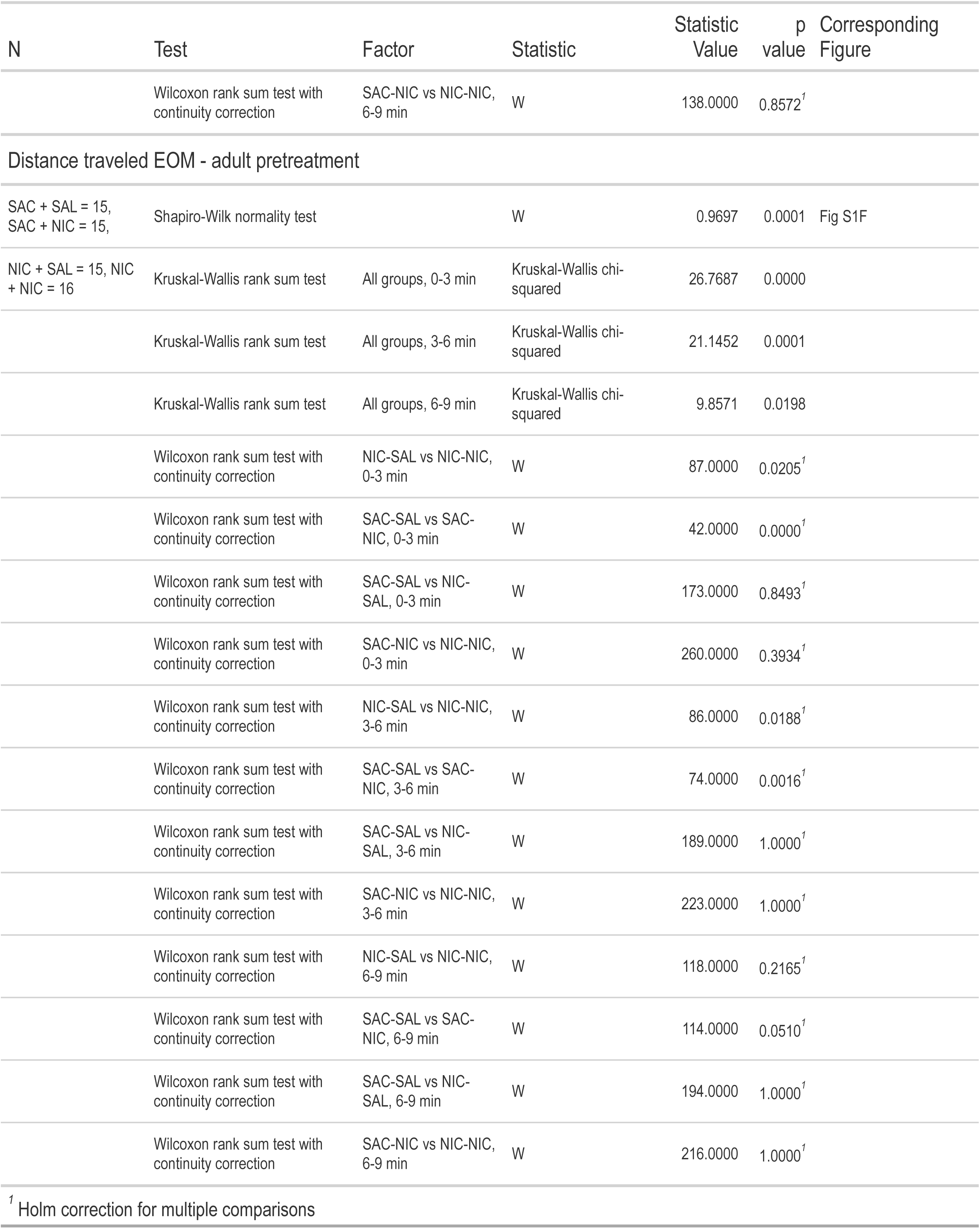
Detailed Statistics for Supplementary Figure 1.

**Table S2.**
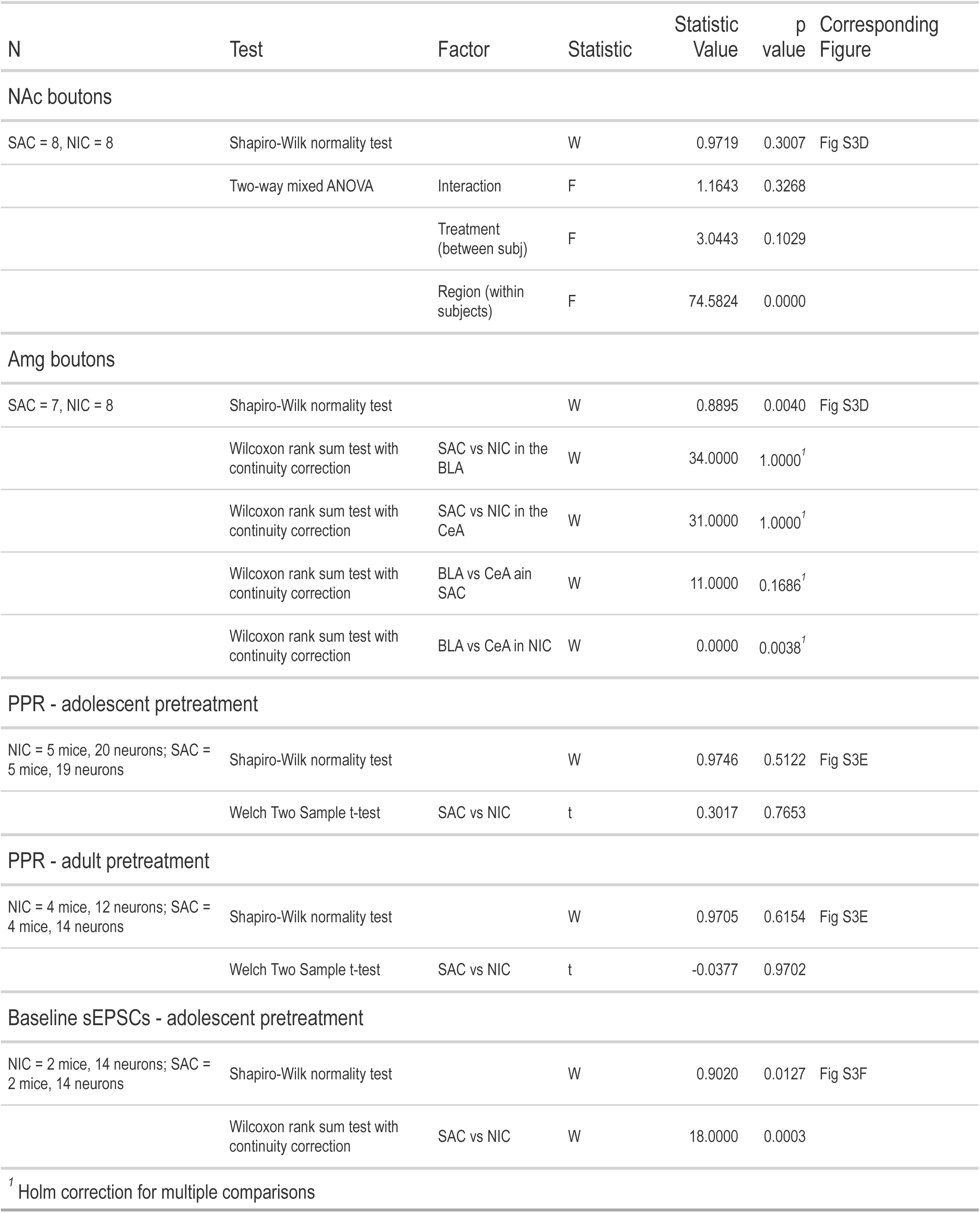
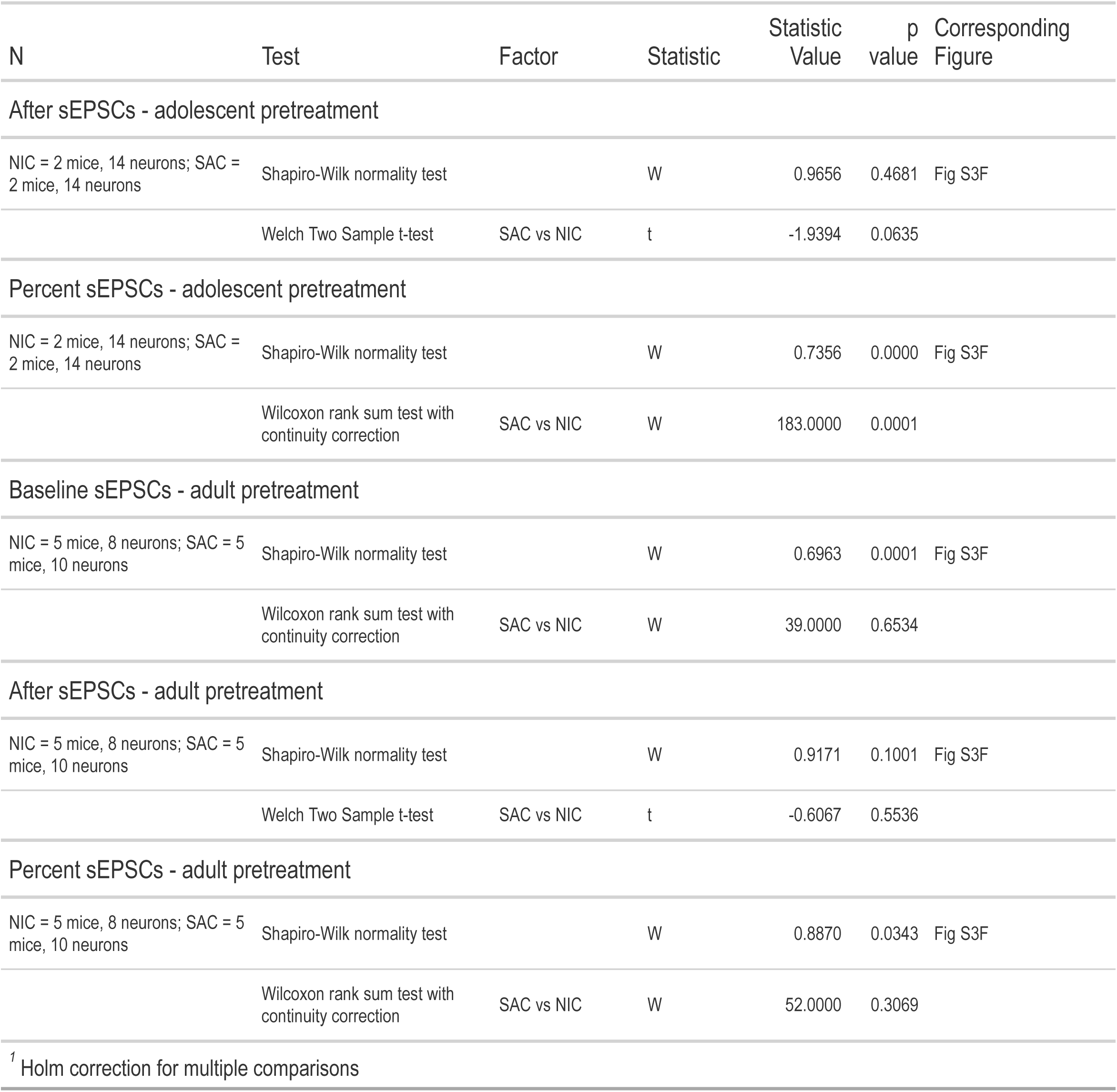
Detailed Statistics for Figure S3.

## Materials and methods

### Animals

All experiments and procedures were performed in accordance with European Commission directives 219/1990, 220/1990 and 2010/63, and approved by Sorbonne Université or the ESPCI. Wild-type (WT) C57BL/6 mice (Janvier Labs, France) or DATiCre mice (from François Tronche^105^) were maintained on a 12-h light–dark cycle (light on at 0800 h) and given ad libitum access to food and water unless noted.

### Drugs

#### Pretreatment regimen

Nicotine tartrate salt (Glentham Biosciences) was dissolved into a 2% Saccharin (Sigma) solution to a final concentration of 100µg/mL and pH was adjusted to 7.2±0.2. Mice had access to either the nicotine solution or a 2% saccharin solution during one week.

#### Experimental solutions

Nicotine tartrate salt was solubilized in a physiological saline solution (0.9% NaCl) with pH adjusted to 7.2±0.2, and was injected intravenously (IV) at a 30 µg/kg dose for juxtacellular recordings, or intra-peritoneally (IP) at a 0.5 mg/kg dose for the elevated O-maze (EOM) test and for the activity mapping experiment. For the Two bottle choice task, Nicotine tartrate salt was dissolved in water to a final concentration of 10µg/mL, 50µg/mL, 100µg/mL, or 200µg/mL and pH was adjusted to 7.2±0.2. All concentrations are expressed as free base.

### Behavior

#### Two-bottle choice

Mice were single-housed in a cage with two drinking bottles where the volume change was measured continuously every minute with an automated acquisition system (TSE system, Germany). Mice were first presented with water in both bottles for a four-day habituation period, and the position of the bottles (i.e. right or left side of the cage) was swapped after 2 days. For each of the test solutions the same pattern was used; mice had access to each of the solutions tested during 4 consecutive days, and the position of the test solution was changed every 2 days (Figure 1A). Mice were first tested for sucrose preference by changing the solution in one bottle to a 0.5% sucrose solution, while the other continued to contain plain water. After 4 days, the test solution was changed to 1% sucrose. Nicotine preference was tested with consecutively increasing doses of nicotine (10, 50, 100 and 200 µg/ml, free base) in the test bottle, the control bottle remained full of plain water. Sweetening the nicotine and control solution with saccharin was avoided as it can obscure the difference in value between the two choices.^63,106–108^ Mice were weighed every other day to quantify the nicotine intake in mg/kg/day. Mice showing a strong side bias (preference <20% or >80% for one side) during the habituation period were removed from the analyses. While measurements took place continuously, sessions were defined as running overnight each test day from 20:00 to 14:00, encompassing the most active drinking periods of the day (Figure 1B) and homogenizing the measurement periods between days with continuous recording and days when the recording had to be stopped in order to change the position and/or contents of the bottles. All solution changes thus occurred during the 14:00-20:00 ‘off’ time window. Minute-by-minute measurements were thresholded at a value of 0.1 ml, with values greater than 0.1ml/min representing less than 0.01% of the full data set and considered to be erroneous measurements (leaking, sensor issue).

#### Elevated O-maze test

All behavioral tests were conducted during the light period of the animal cycle (between 1:00 and 7:00PM). The raw data for behavioral experiments were acquired as video files. The elevated O-maze (EOM) apparatus consists of two open (stressful) and two enclosed (protecting) elevated arms that together form a zero or circle (diameter of 50 cm, height of 58 cm, 10 cm-wide circular platform). Time spent in exploring enclosed versus open arms indicates then the anxiety level of the animal. The test lasts 10 minutes: mice are injected 1 minute before the test, and then put in the EOM for 9 minutes. Mice are placed in an open arm at one of the four entrances of a closed arm, with the entrance (e.g.1-4) distributed pseudorandomly across the experiment. Time spent in open or closed arms was extracted frame-by frame using the open-source video analysis pipeline ezTrack^109^. Mice were habituated to the stress of handling and injection for a minimum of one week before testing.

### Brain clearing and activity mapping

#### Experimental design and perfusion

Mice were injected with i.p. saline or nicotine and kept in a dim, quiet room for one hour before perfusion to minimize off-target cFos expression. Mice were then perfused with 1X PBS followed by 20mL of 4% paraformaldehyde (PFA, Electron Microscopy Services). Brains were carefully dissected from the skull, and stored in PFA overnight. Brains were stored in PBS with 0.01% Sodium Azide (Sigma-Aldrich, Germany) until clearing.

#### iDISCO+ whole brain immunolabeling

Whole brain clearing and immunostaining was performed following the iDISCO+ protocol previously described previously ^66^ with minimal modifications. All the steps of the protocol were done at room temperature with gentle shaking unless otherwise specified. All the buffers were supplemented with 0.01% Sodium Azide (Sigma-Aldrich, Germany) to prevent bacterial and fungal growth. Briefly, perfused brains were dehydrated in an increasing series of methanol (Sigma-Aldrich, France) dilutions in water (washes of 1 hour in methanol 20%, 40%, 60%, 80% and 100%). An additional wash of 2 hours in methanol 100% was done to remove residual water. Once dehydrated, samples were incubated overnight in a solution containing a 66% dichloromethane (Sigma-Aldrich, Germany) in methanol, and then washed twice in methanol 100% (4 hours each wash). Samples were then bleached overnight at 4 C in methanol containing a 5% of hydrogen peroxide (Sigma-Aldrich). Rehydration was done by incubating the samples in methanol 60%, 40% and 20% (1 hour each wash). After methanol pretreatment, samples were washed in PBS twice 15 minutes and 1 hour in PBS containing a 0,2% of Triton X-100 (Sigma-Aldrich) and further permeabilized by a 24 hours incubation at 37C in Permeabilization Solution, composed by 20% dimethyl sulfoxide (Sigma-Aldrich), 2,3% Glycine (Sigma-Aldrich, USA) in PBS-T. In order to start the immunostaining, samples were first blocked with 0,2% gelatin (Sigma-Aldrich) in PBS-T for 24 hours at 37C, the same blocking buffer was used to prepare antibody solutions. Brains were incubated with anti c-Fos primary (Synaptic systems 226-003) for 10 days at 37C with gentle shaking, then washed in PBS-T (twice 1 hour and then overnight), and finally newly incubated for 10 days with secondary antibodies. Secondary antibodies raised in donkeys, conjugated to Alexa 647 were used (Life Technologies). After immunostaining, the samples were washed in PBS-T (twice 1 hour and then overnight), dehydrated in a methanol/water increasing concentration series (20%, 40%, 60%, 80%, 100% one hour each and then methanol 100% overnight), followed by a wash in 66% dichloromethane – 33% methanol for 3 hours. Methanol was washed out with two final washes in dichloromethane 100% (15 min each) and finally the samples were cleared and stored in dibenzyl ether (Sigma-Aldrich) until light sheet imaging.

#### Light sheet microscopy

The acquisitions were done on a LaVision Ultramicroscope II equipped with infinity-corrected objectives. The microscope was installed on an active vibration filtration device, itself put on a marble compressed-air table. Imaging was done with the following filters: 595/40 for Alexa Fluor-555, and -680/30 for Alexa Fluor-647. The microscope was equipped with the following laser lines: OBIS-561nm 100mW, OBIS-639nm 70mW, and used the 2nd generation LaVision beam combiner. The images were acquired with an Andor CMOS sNEO camera. Main acquisitions were done with the LVMI-Fluor 4X/O.3 WD6 LaVision Biotec objective. The microscope was connected to a computer equipped with SSD drives to speed up the acquisition. The brain was positioned in sagittal orientation, cortex side facing the light sheet, to maximize image quality and consistency. A field of view of 1000 x 1300 pixels was cropped at the center of the camera sensor. The light sheet numerical aperture was set to NA-0.03. The 3 light sheets facing the cortex were used, while the other side illumination was deactivated to improve the axial resolution. Beam width was set to the maximum. Laser powers were set to 40-60% (639nm). The center of the light sheet in x was carefully calibrated to the center of the field. z steps were set to 6mm. Tile overlaps were set to 10%. The whole acquisition takes about 1h per hemisphere. At the end of the acquisition, the objective is changed to a MI PLAN 1.1X/0.1 for the reference scan at 488nm excitation (tissue autofluorescence). The field of view is cropped to the size of the brain, and the z-steps are set to 6mm, and light sheet numerical aperture to 0.03 NA. It is important to crop the field of view to the size of the brain for subsequent alignment steps.

#### Computing Resources

The data were automatically transferred every day from the acquisition computer to a Lustre server for storage. The processing with ClearMap was done on local workstations, either Dell Precision T7920 or HP Z840. Each workstation was equipped with 2 Intel Xeon Gold 6128 3.4G 6C/12T CPUs, 512Gb of 2666MHz DDR4 RAM, 4x1Tb NVMe Class 40 Solid State Drives in a RAID0 array (plus a separate system disk), and an NVIDIA Quadro P6000, 24Gb VRAM video card. The workstations were operated by Linux Ubuntu 20.04LTS. ClearMap 2.0 was used on Anaconda Python 3.7 environment.

#### ClearMap Fos+ cell counting

Tiled acquisitions of Fos-immunolabeled iDISCO+ cleared brains scanned with the light sheet microscope were processed with ClearMap 2 to generate both voxel maps of Fos cell densities, as well as region-based statistics of cell counts^67,110^. Briefly, stitched images were processed for background removal, on which local maxima were detected to place initial seeds for the cells. A watershed was done on each seed to estimate the volume of the cell, and the cells were filtered according to their volume to exclude smaller artefactual maxima. The alignment of the brain to the Allen Brain Atlas (March 2017) was based on the acquired autofluorescence image using Elastix (https://elastix.lumc.nl). Filtered cell’s coordinates were transformed to their reference coordinate in the Allen Brain Atlas common coordinate system^111^. For voxel maps, spheres of 375mm diameter were drawn on each filtered cell. P-Value maps of significant differences between groups were generated using Mann-Whitney U test (SciPy implementation). Aligned voxelized datasets from each group of animals were manually inspected to identify the regional overlaps of p-value clusters, and volcano-plots of regional counts where generated.

### Quantitative neuroanatomy of dopamine terminals

#### Tissue preparation

Mice were perfused with 1X PBS followed by 20mL of 4% paraformaldehyde (Sigma). Brains were postfixed in PFA overnight, transferred to PBS before sectioning. Serial 50µm thick coronal sections were made on a Leica vibratome (VTS1000) within one week of perfusion, and stored free-floating in PBS. Free-floating brain sections were then incubated for 1 hour at 4C in a blocking solution of phosphate-buffered saline (PBS) containing 3% bovine serum albumin (BSA, Sigma; A4503) (vol/vol) and 0.2% Triton X-100 (vol/vol), and then incubated overnight at 4C with a sheep anti-tyrosine hydroxylase antibody (anti-TH, Milipore, Ab1542) diluted 1:500 in PBS containing 1.5% BSA and 0.2% Triton X-100. The following day, sections were rinsed with PBS, and then incubated for 3 hours at room temperature with Cy3-conjugated anti-sheep secondary antibody (Jackson ImmunoResearch, 715-165-147) diluted 1:500 in a solution of 1.5% BSA in PBS. After three rinses in PBS, slices were wet-mounted using Prolong Gold Antifade Reagent (Invitrogen, P36930)

#### Confocal image acquisition and deconvolution

Images stacks were taken with a Confocal Laser Scanning Microscope (A1, Nikon) equipped with a 60x 1.4 NA objective (oil immersion, Nikon) with pinhole aperture set to 1 Airy Unit, pixel size of 50 nm and z-step of 200 nm. Excitation wavelength and emission range for Cy-3 labeling was: ex. 561, em. 570-630. Laser intensity was set so that each image occupies the full dynamic range of the detector. Deconvolution using Maximum Likelihood Estimation algorithm was performed with Huygens software (Scientific Volume Imaging).^112^ 150 iterations were applied in classical mode, background intensity was averaged from the voxels with lowest intensity, and signal to noise ratio values were set to a value of 20.

#### Segmentation of presynaptic boutons from confocal images

Segmentation of TH+ boutons was performed in 3D in FIJI (fiji.net) using a procedure based on image analysis tools developed in the ImageJ plugin 3DImageSuite.^113– 115^ First local maxima were detected. Histogram analysis and image inspection allowed the definition of a threshold intensity so that only local maxima from presynaptic objects were retained. The local maxima were then used as seeds around which 3D intensity distribution was fitted to a Gaussian curve. The intensity value which defines 95% of the area of the gauss curve was chosen as threshold for the border of the object. Buttons of similar size but different intensity are thus extracted as object of similar size. The border of the object was determined following the so-called block algorithm; starting around the local maxima, voxels which followed three criteria were included in the object (intensity above the threshold, intensity lower than previously included voxel, and the inclusion is validated if neighboring voxels are included as well). Counts were used to determine a density for the obtained image stack and normalized to a standard volume of 10µm x 10µm x 10µm to make comparisons between images with different stack sizes, then exported to .csv.

### In vivo electrophysiology

Mice were deeply anaesthetized with (1) an IP injection of chloral hydrate (400 mg/kg), supplemented as required to maintain optimal anesthesia throughout the experiment, or (2) isoflurane delivered continuously (5% induction, 2-3% maintenance; TEMSega). The scalp was opened and a hole was drilled in the skull above the location of the VTA. Intravenous administration of saline or nicotine (30µg/kg) was carried out through a catheter (30G needle connected to polyethylene tubing PE10) connected to a Hamilton syringe, into the saphenous vein of the animal. Extracellular recording electrodes were constructed from 1.5 mm outer diameter / 1.17 mm inner diameter borosilicate glass tubing (Harvard Apparatus) using a vertical electrode puller (Narishige). The tip was broken straight and clean under microscopic control to obtain a diameter of about 1 µm. The electrodes were filled with a 0.5% NaCl solution containing 1.5% of neurobiotin^®^ tracer (VECTOR laboratories) yielding impedances of 6-9 MΩ. Electrical signals were amplified by a high-impedance amplifier (Axon Instruments) and monitored audibly through an audio monitor (A.M. Systems Inc.). The signal was digitized, sampled at 25 kHz, and recorded on a computer using Spike2 software (Cambridge Electronic Design) for later analysis. The electrophysiological activity was sampled in the central region of the VTA (coordinates: between 3.1 to 4 mm posterior to bregma, 0.3 to 0.7 mm lateral to midline, and 4 to 4.8 mm below brain surface). Individual electrode tracks were separated from one another by at least 0.1 mm in the horizontal plane. Spontaneously active DA neurons were identified based on previously established electrophysiological criteria^116^.

After recording, a subset of nicotine-responsive cells were labelled by electroporation of their membrane: successive currents squares were applied until the membrane breakage, to fill the cell soma with neurobiotin contained into the glass pipet^117^. To be able to establish correspondence between neurons responses and their localization in the VTA, we labeled one type of response per mouse: solely activated neurons or solely inhibited neurons, with a limited number of cells per brain (1 to 4 neurons maximum, 2 by hemisphere), always with the same concern of localization of neurons in the VTA.

### Ex vivo patch-clamp recordings

Mice were anesthetized (Ketamine 150 mg/kg; Xylazine 10 mg/kg) and transcardially perfused with aCSF for slice preparation. For VTA recordings, horizontal 250 μm slices were obtained in bubbled ice-cold 95% O2/5% CO2 aCSF containing (in mM): KCl 2.5, NaH2PO4 1.25, MgSO4 10, CaCl2 0.5, glucose 11, sucrose 234, NaHCO3 26. Slices were then incubated in aCSF containing (in mM): NaCl 119, KCl 2.5, NaH2PO4 1.25, MgSO4 1.3, CaCl2 2.5, NaHCO3 26, glucose 11, at 37 °C for 1 h, and then kept at room temperature. Slices were transferred and kept at 32–34 °C in a recording chamber superfused with 2.5 ml/min aCSF. Visualized whole-cell voltage-clamp recording technique was used to measure synaptic responses using an upright microscope (Olympus France). Putative DA neurons were recorded in the lateral VTA and identified using criteria such as localization and cell body size, as well as electrophysiological signature (e.g., broad action potential, and large Ih current).^116^ Spontaneous excitatory postsynaptic currents (sEPSCs) induced by puffing nicotine onto DA cells were measured, as well as analyses of paired-pulse ratios (PPR) and AMPA-R/NMDA-R ratios as an index of synaptic adaptations.

Experiments were obtained using a Multiclamp 700B (Molecular Devices, Sunnyvale, CA). Signals were collected and stored using a Digidata 1440 A converter and pCLAMP 10.2 software (Molecular Devices, CA). In all cases, analyses were performed using Clampfit 10.2 (Axon Instruments, USA) and Prism (Graphpad, USA). Paired-pulse ratios (PPR) and AMPA-R/NMDA-R ratios were assessed in voltage-clamp mode using an internal solution containing (in mM) 130 CsCl, 4 NaCl, 2 MgCl2, 1.1 EGTA, 5 HEPES, 2 Na2ATP, 5 sodium creatine phosphate, 0.6 Na3GTP, and 0.1 spermine. The PPR protocol consisted in two evoked pulses 50 ms apart applied every 15 s at V= -60 mV. Synaptic currents were evoked by stimuli (10 µs) at 0.15 Hz through a glass pipette placed 200 µm from the patched neurons. Voltage was then raised to V= + 40 mV and synaptic currents were evoked in the absence and in the presence of AMPA-R antagonist DNQX as previously described for AMPA-R/NMDA-R ratios.^118^

For analyses of peak amplitudes of nAChR currents and nicotinic modulation of sEPSCs, nicotine was applied by a ‘puff’ with air-pressure pulses controlled by a Picospritzer III (General Valve). A drug-filled pipette was moved within 20-40µm from the recorded neuron and a pClamp protocol triggered a puff application of nicotine (10 mM) onto the recorded neuron with a 20 – 80 ms, 20-psi pressure ejection. Peak amplitude (pA) was compared between groups. To compare the frequency and amplitudes of sEPSCs in response to acute NIC exposure between treated animals and controls, epochs of at least 30 s were recorded before and after puff application of nicotine on VTA DA neurons. Analyses were performed using Clampfit 10.2 (Axon Instruments, USA) and Prism (Graphpad, USA).

### DREADD experiments

DAT-Cre mice, in which Cre recombinase expression is restricted to DA neurons without disrupting endogenous dopamine transporter (DAT) expression,^105^ were weaned at PND 21±1 and treated with nicotine (100µg/mL in 2% saccharine) for one week. When they reached adulthood, they were injected with Cre-dependent (DIO) DREADD or control (tdTomato) viruses. Adult DAT-Cre mice were anesthetized with a mixture of oxygen (1 L/min) and 3-4% isoflurane (Vetflurane, Virbac) for the induction of anesthesia, and then placed on a warming pad in a stereotaxic frame (David Kopf) and maintained under anesthesia throughout the surgery at 2% isoflurane. A local anesthetic (Lurocaine) was applied at the location of the scalp incision and 0.1 µL of buprenorphine (Buprecare, 1 mg/kg) was injected subcutaneously before the procedure. Mice then received bilateral injections of a retrogradely-transported inhibitory DREADD AAV (ssAAV-retro/2-hSyn1-dlox-hM4D(Gi)_mCherry(rev)-dlox-WPRE-hGHp(A), Viral Vector Facility Zurich) or control virus (ssAAV-retro/2-hEF1a-dlox-dTomato-EGFP(rev)-dlox-WPRE-hGHp(A), Viral Vector Facility Zurich) into the medial NAc (bregma +1.7 mm, lateral ±0.45 mm, ventral -4.1 mm).

Mice were tested in the EOM 4-5 weeks after viral injections to allow full expression of the DREADD or control viruses at the level of the VTA. Clozapine-N-oxide (5 mg/kg, CNO) was injected i.p. one hour before the mice received nicotine or saline and entered into the EOM task, as described above. While CNO can have off-target effects^119^, the 5mg/kg dose used in our experiments is below the threshold identified for off-target effects. In support of this idea, we did not see any differences in saline-injected mice under CNO, regardless of their virus (DREADD or control) status.

Following the EOM, mice were perfused with 1X PBS followed by 20mL of 4% paraformaldehyde (Sigma). Brains were postfixed in PFA overnight, and transferred to PBS before sectioning. Serial 50µm thick coronal sections were made on a Leica vibratome (VTS1000) within one week of perfusion, and stored free-floating in PBS. Free-floating brain sections were then incubated for 1 hour at 4C in a blocking solution of phosphate-buffered saline (PBS) containing 3% bovine serum albumin (BSA, Sigma; A4503) (vol/vol) and 0.2% Triton X-100 (vol/vol), and then incubated overnight at 4C with a sheep anti-tyrosine hydroxylase antibody (anti-TH, Milipore, Ab1542) diluted 1:500 in PBS containing 1.5% BSA and 0.2% Triton X-100. The following day, sections were rinsed with PBS, and then incubated for 3 hours at room temperature with Cy5-conjugated anti-sheep secondary antibody (Jackson ImmunoResearch, 713-175-147) diluted 1:500 in a solution of 1.5% BSA in PBS. After three rinses in PBS, slices were wet-mounted using Prolong Gold Antifade Reagent containing DAPI (Invitrogen, P36931). Viral expression was evaluated by observing the unamplified expression of mCherry or tdTomato via a Rhodamine filter on an epifluorescent microscope (Zeiss Axio Imager), dopamine neurons were identified by TH staining via a Cy5 filter, and non-dopamine cell bodies were evaluated by DAPI staining. All neurons expressing the DREADD or control viruses co-expressed TH, and staining was limited to the medial VTA in line with previous reports on the location of neurons projecting to the medial NAc.^43^

### Statistical analysis

All statistical analysis and graphs were made using R, a language and environment for statistical computing (Team, 2005, http://www.r-project.org), with the exception of in vitro electrophysiology data, which was analyzed in Prism (GraphPad). Data from behavioral experiments was exported from acquisition software (TSE system) or from ezTrack video tracking as a .csv file, and read directly into R. Quantifications from ClearMap or from bouton analysis in FIJI were likewise exported as .csv files and read directly into R. CSV files of cell counts by region were imported into R from ClearMap. Correlations and clustering were then performed in R using the {stats} package, and networks were created {igraph} package. For the measurement of neuronal activity in single-unit recordings, timestamps of action potentials were extracted in Spike 2, analyzed in R, and expressed as the average firing frequency (in Hz) and the percentage of spikes-within-burst (%SWB = number of spikes within burst divided by total number of spikes in a given window). Neuronal basal activity was defined on recordings of a minimum of three minutes. Firing frequency was quantified on overlapping 60-second windows shifted by 15-second time steps. For each neuron, the firing frequency was rescaled as a percentage of its baseline value averaged during 3 minutes before nicotine injection. The responses to nicotine are thus presented as a percentage of variation from baseline (mean ± S.E.M.). The effect of nicotine was assessed by comparison of the maximum of firing frequency variation induced by nicotine and saline injection. For activated (respectively inhibited) neurons, the maximal (respectively minimal) value of the firing frequency was measured within the response period (3 minutes) that followed nicotine or saline injection. The results are presented as mean ± S.E.M. of the difference of maximum variation after nicotine or saline.

## Statistics and Reproducibility

All experiments were replicated with success.

## Notes

### Competing Interest Statement

The authors have declared no competing interest.

## References

1. Ezzati, M. et al. Selected major risk factors and global and regional burden of disease. Lancet 360, 1347–1360(2002).

2. SAMHSA. Results from the 2011 National Survey on Drug Use and Health: Summary of national findings. NSDUH Series H-44, HHS Publication No. (SMA) 12-4713. Rockville, MD: Substance Abuse and Mental Health Services Administration (2012).

3. Anthony, J. C. & Petronis, K. R. Early-onset drug use and risk of later drug problems. Drug Alcohol Depen 40, 9–15 (1995).

4. Grant, B. F. & Dawson, D. A. Age of onset of drug use and its association with DSM-IV drug abuse and dependence: results from the National Longitudinal Alcohol Epidemiologic Survey. Journal of substance abuse 10, 163 173 (2003).

5. Grant, B. F. Age at smoking onset and its association with alcohol consumption and DSM-IV alcohol abuse and dependence: Results from the national longitudinal alcohol epidemiologic survey. J Subst Abuse 10, 59–73 (1998).

6. Jamal, M., Does, A. J. W. V. D, Penninx, B. W. J. H. & Cuijpers, P. Age at Smoking Onset and the Onset of Depression and Anxiety Disorders. Nicotine Tob Res 13, 809–819 (2011).

7. Choi, W. S., Patten, C. A., Gillin, J. C., Kaplan, R. M. & Pierce, J. P. Cigarette smoking predicts development of depressive symptoms among U.S. Adolescents. Ann Behav Med 19, 42–50 (1997).

8. Kota, D., Martin, B. R., Robinson, S. E. & Damaj, M. I. Nicotine Dependence and Reward Differ between Adolescent and Adult Male Mice. J. Pharmacol. Exp. Ther. 322, 399–407 (2007).

9. Shram, M. J., Funk, D., Li, Z. & Lê, A. D. Periadolescent and adult rats respond differently in tests measuring the rewarding and aversive effects of nicotine. Psychopharmacology 186, 201 (2006).

10. Belluzzi, J. D., Lee, A. G., Oliff, H. S. & Leslie, F. M. Age-dependent effects of nicotine on locomotor activity and conditioned place preference in rats. Psychopharmacology 174, 389 395 (2004).

11. Adriani, W., Macrì, S., Pacifici, R. & Laviola, G. Peculiar Vulnerability to Nicotine Oral Self-administration in Mice during Early Adolescence. Neuropsychopharmacology 27, 212–224 (2002).

12. Faraday, M. M., Elliott, B. M. & Grunberg, N. E. Adult vs. adolescent rats differ in biobehavioral responses to chronic nicotine administration. Pharmacol. Biochem. Behav. 70, 475–489 (2001).

13. Schassburger, R. L. et al. Adolescent Rats Self-Administer Less Nicotine Than Adults at Low Doses. Nicotine Tob. Res. 18, 1861–1868 (2016).

14. Vastola, B. J., Douglas, L. A., Varlinskaya, E. I. & Spear, L. P. Nicotine-induced conditioned place preference in adolescent and adult rats. Physiol. Behav. 77, 107–114 (2002).

15. Slotkin, T. A., Bodwell, B. E., Ryde, I. T. & Seidler, F. J. Adolescent nicotine treatment changes the response of acetylcholine systems to subsequent nicotine administration in adulthood. Brain Res. Bull. 76, 152– 165 (2008).

16. Kota, D., Robinson, S. E. & Damaj, M. I. Enhanced nicotine reward in adulthood after exposure to nicotine during early adolescence in mice. Biochem. Pharmacol. 78, 873–879 (2009).

17. Bracken, A. L., Chambers, R. A., Berg, S. A., Rodd, Z. A. & McBride, W. J. Nicotine exposure during adolescence enhances behavioral sensitivity to nicotine during adulthood in Wistar rats. Pharmacol. Biochem. Behav. 99, 87–93 (2011).

18. Chellian, R. et al. Adolescent nicotine and tobacco smoke exposure enhances nicotine self-administration in female rats. Neuropharmacology 108243 (2020) doi:10.1016/j.neuropharm.2020.108243.

19. Kallupi, M., Guglielmo, G. de, Larrosa, E. & George, O. Exposure to passive nicotine vapor in male adolescent rats produces a withdrawal-like state and facilitates nicotine self-administration during adulthood. Eur Neuropsychopharm (2019) doi:10.1016/j.euroneuro.2019.08.299.

20. Adriani, W., Deroche-Gamonet, V., Moal, M. L., Laviola, G. & Piazza, P. V. Preexposure during or following adolescence differently affects nicotine-rewarding properties in adult rats. Psychopharmacology 184, 382 390 (2006).

21. Natividad, L. A., Torres, O. V., Friedman, T. C. & O’Dell, L. E. Adolescence is a period of development characterized by short- and long-term vulnerability to the rewarding effects of nicotine and reduced sensitivity to the anorectic effects of this drug. Behav Brain Res 257C, 275 285 (2013).

22. Leslie, F. M. Unique, long-term effects of nicotine on adolescent brain. Pharmacol Biochem Be 173010 (2020) doi:10.1016/j.pbb.2020.173010.

23. Thorpe, H. H. A., Hamidullah, S., Jenkins, B. W. & Khokhar, J. Y. Adolescent Neurodevelopment and Substance Use: Receptor Expression and Behavioral Consequences. Pharmacol Therapeut 107431 (2019) doi:10.1016/j.pharmthera.2019.107431.

24. Abreu-Villaça, Y. et al. Short-term adolescent nicotine exposure has immediate and persistent effects on cholinergic systems: critical periods, patterns of exposure, dose thresholds. Neuropsychopharmacol 28, 1935–1949 (2003).

25. Trauth, J. A., Seidler, F. J., McCook, E. C. & Slotkin, T. A. Adolescent nicotine exposure causes persistent upregulation of nicotinic cholinergic receptors in rat brain regions. Brain Res 851, 9–19 (1999).

26. Trauth, J. A., McCook, E. C., Seidler, F. J. & Slotkin, T. A. Modeling adolescent nicotine exposure: effects on cholinergic systems in rat brain regions. Brain Res 873, 18–25 (2000).

27. Doura, M. B., Luu, T. V., Lee, N. H. & Perry, D. C. Persistent gene expression changes in ventral tegmental area of adolescent but not adult rats in response to chronic nicotine. Neuroscience 170, 503–513 (2010).

28. Somerville, L. H. & Casey, B. Developmental neurobiology of cognitive control and motivational systems. Curr Opin Neurobiol 20, 236–241 (2010).

29. Casey, B. J. Beyond Simple Models of Self-Control to Circuit-Based Accounts of Adolescent Behavior. Annu Rev Psychol 66, 1–25 (2015).

30. Nussenbaum, K. & Hartley, C. A. Reinforcement learning across development: What insights can we draw from a decade of research? Dev Cogn Neuros-neth 40, 100733 (2019).

31. Barth, B., Portella, A. K., Dubé, L., Meaney, M. J. & Silveira, P. P. Early Life Origins of Ageing and Longevity. in Early Life Origins of Ageing and Longevity 121–140 (2019). doi:10.1007/978-3-030-24958-8_7.

32. Reynolds, L. M. & Flores, C. Adolescent dopamine development: connecting experience with vulnerability or resilience to psychiatric disease. in Diagnosis, Management and Modeling of Neurodevelopmental Disorders (eds. Martin, C. R., Preedy, V. R. & Rajendram, R.) (Academic Press, 2021). doi:10.1016/b978-0-12-817988-8.00026-9.

33. Reynolds, L. M. & Flores, C. Mesocorticolimbic Dopamine Pathways Across Adolescence: Diversity in Development. Front Neural Circuit 15, 735625 (2021).

34. Changeux, J.-P. Nicotine addiction and nicotinic receptors: lessons from genetically modified mice. Nat Rev Neurosci 11, 389 401 (2010).

35. Mameli-Engvall, M. et al. Hierarchical control of dopamine neuron-firing patterns by nicotinic receptors. Neuron 50, 911 921 (2006).

36. Maskos, U. et al. Nicotine reinforcement and cognition restored by targeted expression of nicotinic receptors. Nature 436, 103 107 (2005).

37. Morel, C., Montgomery, S. & Han, M.-H. Nicotine and alcohol: the role of midbrain dopaminergic neurons in drug reinforcement. Eur J Neurosci 114, 13012 (2018).

38. Faure, P., Tolu, S., Valverde, S. & Naudé, J. Role of nicotinic acetylcholine receptors in regulating dopamine neuron activity. Neuroscience 282, 86 100 (2014).

39. Wills, L. et al. Neurobiological Mechanisms of Nicotine Reward and Aversion. Pharmacol. Rev. 74, 271–310 (2022).

40. Placzek, A. N., Zhang, T. A. & Dani, J. A. Age dependent nicotinic influences over dopamine neuron synaptic plasticity. Biochem Pharmacol 78, 686 692 (2009).

41. Jobson, C. L. M. et al. Adolescent Nicotine Exposure Induces Dysregulation of Mesocorticolimbic Activity States and Depressive and Anxiety-like Prefrontal Cortical Molecular Phenotypes Persisting into Adulthood. Cereb Cortex 29, (2018).

42. Lammel, S., Ion, D. I., Roeper, J. & Malenka, R. C. Projection-specific modulation of dopamine neuron synapses by aversive and rewarding stimuli. Neuron 70, 855 862 (2011).

43. Lammel, S. et al. Unique properties of mesoprefrontal neurons within a dual mesocorticolimbic dopamine system. Neuron 57, 760 773 (2008).

44. Phillips, R. A. et al. An atlas of transcriptionally defined cell populations in the rat ventral tegmental area. Cell Reports 39, 110616 (2022).

45. Poulin, J.-F. et al. Mapping projections of molecularly defined dopamine neuron subtypes using intersectional genetic approaches. Nat Neurosci 21, 1 (2018).

46. Margolis, E. B., Hjelmstad, G. O., Fujita, W. & Fields, H. L. Direct Bidirectional µ-Opioid Control of Midbrain Dopamine Neurons. J Neurosci 34, 14707 14716 (2014).

47. Juarez, B. & Han, M.-H. Diversity of Dopaminergic Neural Circuits in Response to Drug Exposure. Neuropsychopharmacol 41, (2016).

48. Lammel, S., Lim, B. K. & Malenka, R. C. Reward and aversion in a heterogeneous midbrain dopamine system. Neuropharmacology 76 Pt B, 351–359 (2014).

49. Eddine, R. et al. A concurrent excitation and inhibition of dopaminergic subpopulations in response to nicotine. Sci Rep-uk 5, 8184 (2015).

50. Liu, C. et al. An inhibitory brainstem input to dopamine neurons encodes nicotine aversion. Neuron (2022) doi:10.1016/j.neuron.2022.07.003.

51. Kutlu, M. G. & Gould, T. J. Nicotine modulation of fear memories and anxiety: Implications for learning and anxiety disorders. Biochem Pharmacol 97, 498–511 (2015).

52. Picciotto, M. R. & Mineur, Y. S. Molecules and circuits involved in nicotine addiction: The many faces of smoking. Neuropharmacology 76, 545–553 (2014).

53. Nguyen, C. et al. Nicotine inhibits the VTA-to-amygdala dopamine pathway to promote anxiety. Neuron (2021) doi:10.1016/j.neuron.2021.06.013.

54. Morel, C. et al. Midbrain projection to the basolateral amygdala encodes anxiety-like but not depression-like behaviors. Nat Commun 13, 1532 (2022).

55. Reynolds, L. M. et al. Amphetamine disrupts dopamine axon growth in adolescence by a sex-specific mechanism in mice. Nat. Commun. 14, 4035 (2023).

56. Reynolds, L. M. et al. Amphetamine in Adolescence Disrupts the Development of Medial Prefrontal Cortex Dopamine Connectivity in a dcc-Dependent Manner. Neuropsychopharmacol 40, 1101–1112(2015).

57. Reynolds, L. M. et al. Early Adolescence is a Critical Period for the Maturation of Inhibitory Behavior. Cereb Cortex 29, 3676–3686 (2019).

58. Glantz, S., Jeffers, A. & Winickoff, J. P. Nicotine Addiction and Intensity of e-Cigarette Use by Adolescents in the US, 2014 to 2021. Jama Netw Open 5, e2240671 (2022).

59. Riggs, N., Chou, C.-P., Li, C. & Pentz, M. A. Adolescent to emerging adulthood smoking trajectories: When do smoking trajectories diverge, and do they predict early adulthood nicotine dependence? Nicotine Tob. Res. 9, 1147–1154 (2007).

60. Fowler, C. D. & Kenny, P. J. Intravenous nicotine self-administration and cue-induced reinstatement in mice: Effects of nicotine dose, rate of drug infusion and prior instrumental training. Neuropharmacology 61, 687–698 (2011).

61. Mondoloni, S. et al. Prolonged nicotine exposure reduces aversion to the drug in mice by altering nicotinic transmission in the interpeduncular nucleus. eLife 12, (2023).

62. Fowler, C. D., Lu, Q., Johnson, P. M., Marks, M. J. & Kenny, P. J. Habenular α5 nicotinic receptor subunit signalling controls nicotine intake. Nature 471, 597–601 (2011).

63. Bagdas, D. et al. Assessing nicotine dependence using an oral nicotine free-choice paradigm in mice. Neuropharmacology 157, 107669 (2019).

64. Isiegas, C., Mague, S. D. & Blendy, J. A. Sex differences in response to nicotine in C57Bl/6:129SvEv mice. Nicotine Tob Res 11, 851–858 (2009).

65. Robinson, S. F., Marks, M. J. & Collins, A. C. Inbred mouse strains vary in oral self-selection of nicotine. Psychopharmacology 124, 332–339 (1996).

66. Renier, N., et al. iDISCO: A Simple, Rapid Method to Immunolabel Large Tissue Samples for Volume Imaging. Cell 159, 896–910 (2014).

67. Renier, N. et al. Mapping of Brain Activity by Automated Volume Analysis of Immediate Early Genes. Cell 165, 1789–1802 (2016).

68. Hale, M. W. et al. Exposure to an open-field arena increases c-Fos expression in a distributed anxiety-related system projecting to the basolateral amygdaloid complex. Neuroscience 155, 659–672 (2008).

69. Singewald, N., Salchner, P. & Sharp, T. Induction of c-Fos expression in specific areas of the fear circuitry in rat forebrain by anxiogenic drugs. Biol Psychiat 53, 275–283 (2003).

70. Singewald, N. & Sharp, T. Neuroanatomical targets of anxiogenic drugs in the hindbrain as revealed by Fos immunocytochemistry. Neuroscience 98, 759–770 (2000).

71. Pich, E. M., Chiamulera, C. & Tessari, M. Neural substrate of nicotine addiction as defined by functional brain maps of gene expression. J Physiology-paris 92, 225–228 (1998).

72. Pagliusi, S. R., Tessari, M., DeVevey, S., Chiamulera, C. & Pich, E. M. The Reinforcing Properties of Nicotine are Associated with a Specific Patterning of c-fos Expression in the Rat Brain. Eur J Neurosci 8, 2247– 2256 (1996).

73. Bellone, C., Mameli, M. & Lüscher, C. In utero exposure to cocaine delays postnatal synaptic maturation of glutamatergic transmission in the VTA. Nat Neurosci 14, 1439–1446(2011).

74. Bellone, C. & Nicoll, R. A. Rapid Bidirectional Switching of Synaptic NMDA Receptors. Neuron 55, 779–785 (2007).

75. Basilico, B. et al. Microglia shape presynaptic properties at developing glutamatergic synapses. Glia 67, 53– 67 (2019).

76. Shram, M. J. & Lê, A. D. Adolescent male Wistar rats are more responsive than adult rats to the conditioned rewarding effects of intravenously administered nicotine in the place conditioning procedure. Behav. Brain Res. 206, 240–244 (2010).

77. Elliott, B. M., Faraday, M. M., Phillips, J. M. & Grunberg, N. E. Effects of nicotine on elevated plus maze and locomotor activity in male and female adolescent and adult rats. Pharmacol. Biochem. Behav. 77, 21–28 (2004).

78. Bariselli, S. et al. Role of VTA dopamine neurons and neuroligin 3 in sociability traits related to nonfamiliar conspecific interaction. Nat Commun 9, 3173 (2018).

79. Miech, R., Johnston, L., O’Malley, P. M., Bachman, J. G. & Patrick, M. E. Adolescent Vaping and Nicotine Use in 2017-2018 - U.S. National Estimates. New Engl J Med 380, 192 193 (2019).

80. Johnson, S. W. & North, R. A. Two types of neurone in the rat ventral tegmental area and their synaptic inputs. J. Physiol. 450, 455–468 (1992).

81. Olson, V. G. & Nestler, E. J. Topographical organization of GABAergic neurons within the ventral tegmental area of the rat. Synapse 61, 87–95 (2007).

82. Yamaguchi, T., Sheen, W. & Morales, M. Glutamatergic neurons are present in the rat ventral tegmental area. Eur. J. Neurosci. 25, 106–118 (2007).

83. Beier, K. T. et al. Circuit Architecture of VTA Dopamine Neurons Revealed by Systematic Input-Output Mapping. Cell 162, 622 634 (2015).

84. Lammel, S. et al. Input-specific control of reward and aversion in the ventral tegmental area. Nature 491, 212 217 (2012).

85. Ford, C. P., Mark, G. P. & Williams, J. T. Properties and Opioid Inhibition of Mesolimbic Dopamine Neurons Vary according to Target Location. J Neurosci 26, 2788–2797 (2006).

86. Mejias-Aponte, C. A., Ye, C., Bonci, A., Kiyatkin, E. A. & Morales, M. A subpopulation of neurochemically-identified ventral tegmental area dopamine neurons is excited by intravenous cocaine. J Neurosci 35, 1965–1978 (2015).

87. Simmons, S. C., Wheeler, K. & Mazei-Robison, M. S. Determination of circuit-specific morphological adaptations in ventral tegmental area dopamine neurons by chronic morphine. Mol Brain 12, 10 (2019).

88. Dannenhoffer, C. A. & Spear, L. P. Age differences in conditioned place preferences and taste aversions to nicotine. Dev. Psychobiol. 58, 660–666 (2016).

89. Wilmouth, C. E. & Spear, L. P. Adolescent and Adult Rats’ Aversion to Flavors Previously Paired with Nicotine. Ann. N. York Acad. Sci. 1021, 462–464 (2004).

90. Wills, L. & Kenny, P. J. Addiction-related neuroadaptations following chronic nicotine exposure. J Neurochem (2021) doi:10.1111/jnc.15356.

91. Fowler, C. D. & Kenny, P. J. Nicotine aversion: Neurobiological mechanisms and relevance to tobacco dependence vulnerability. Neuropharmacology 76, 533–544 (2014).

92. Laviolette, S. R. & Kooy, D. van der. The neurobiology of nicotine addiction: bridging the gap from molecules to behaviour. Nat Rev Neurosci 5, 55–65 (2004).

93. Lidov, H. G. W., Grzanna, R. & Molliver, M. E. The serotonin innervation of the cerebral cortex in the rat—an immunohistochemical analysis. Neuroscience 5, 207–227 (1980).

94. Levitt, P. & Moore, R. Y. Development of the noradrenergic innervation of neocortex. Brain research 162, 243 259 (1979).

95. Suri, D., Teixeira, C. M., Cagliostro, M. K. C., Mahadevia, D. & Ansorge, M. S. Monoamine-sensitive developmental periods impacting adult emotional and cognitive behaviors. Neuropsychopharmacol 40, 88 112 (2014).

96. Vassilev, P. et al. Unique effects of social defeat stress in adolescent male mice on the Netrin-1/DCC pathway, prefrontal cortex dopamine and cognition (Social stress in adolescent vs. adult male mice). Eneuro ENEURO.0045-21.2021 (2021) doi:10.1523/eneuro.0045-21.2021.

97. Baarendse, P. J. J., Counotte, D. S., O’Donnell, P. & Vanderschuren, L. J. M. J. Early social experience is critical for the development of cognitive control and dopamine modulation of prefrontal cortex function. Neuropsychopharmacol 38, 1485–1494 (2013).

98. Jordan, C. J. & Andersen, S. L. Sensitive periods of substance abuse: Early risk for the transition to dependence. Dev Cogn Neuros-neth 25, 29 44 (2017).

99. Lin, W. C. et al. Transient food insecurity during the juvenile-adolescent period affects adult weight, cognitive flexibility, and dopamine neurobiology. Curr. Biol. 32, 3690–3703.e5 (2022).

100. Hoops, D., Reynolds, L. M., Restrepo-Lozano, J. M. & Flores, C. Dopamine Development in the Mouse Orbital Prefrontal Cortex Is Protracted and Sensitive to Amphetamine in Adolescence. Eneuro 5, ENEURO.0372-17.2017 (2018).

101. Walker, D. M. et al. Adolescence and Reward: Making Sense of Neural and Behavioral Changes Amid the Chaos. J Neurosci 37, 10855 10866 (2017).

102. Cross, S. J., Linker, K. E. & Leslie, F. M. Sex-dependent effects of nicotine on the developing brain. J Neurosci Res 95, 422 436 (2017).

103. Okoli, C., Greaves, L. & Fagyas, V. Sex differences in smoking initiation among children and adolescents. Public Heal. 127, 3–10 (2013).

104. Duan, Z., Wang, Y. & Huang, J. Sex Difference in the Association between Electronic Cigarette Use and Subsequent Cigarette Smoking among U.S. Adolescents: Findings from the PATH Study Waves 1–4. Int. J. Environ. Res. Public Heal. 18, 1695 (2021).

105. Turiault, M. et al. Analysis of dopamine transporter gene expression pattern -- generation of DAT-iCre transgenic mice. Febs J 274, 3568–3577(2007).

106. Klein, L. C., Stine, M. M., Vandenbergh, D. J., Whetzel, C. A. & Kamens, H. M. Sex differences in voluntary oral nicotine consumption by adolescent mice: a dose-response experiment. Pharmacol Biochem Be 78, 13 25 (2004).

107. Nesil, T., Kanit, L., Collins, A. C. & Pogun, S. Individual differences in oral nicotine intake in rats. Neuropharmacology 61, 189–201 (2011).

108. Locklear, L. L., McDonald, C. G., Smith, R. F. & Fryxell, K. J. Adult mice voluntarily progress to nicotine dependence in an oral self-selection assay. Neuropharmacology 63, 582–92 (2012).

109. Pennington, Z. T. et al. ezTrack: An open-source video analysis pipeline for the investigation of animal behavior. Sci Rep-uk 9, 19979 (2019).

110. Kirst, C. et al. Mapping the Fine-Scale Organization and Plasticity of the Brain Vasculature. Cell 180, 780–795.e25 (2020).

111. Wang, Q. et al. The Allen Mouse Brain Common Coordinate Framework: A 3D Reference Atlas. Cell 181, 936–953.e20 (2020).

112. Heck, N., Betuing, S., Vanhoutte, P. & Caboche, J. A deconvolution method to improve automated 3D-analysis of dendritic spines: application to a mouse model of Huntington’s disease. Brain Struct Funct 217, 421– 34 (2011).

113. Heck, N. et al. A new automated 3D detection of synaptic contacts reveals the formation of cortico-striatal synapses upon cocaine treatment in vivo. Brain Struct Funct 220, 2953–2966 (2015).

114. Gilles, J.-F., Santos, M. D., Boudier, T., Bolte, S. & Heck, N. DiAna, an ImageJ tool for object-based 3D co-localization and distance analysis. Methods 115, 55–64 (2017).

115. Santos, M. D. et al. Cocaine increases dopaminergic connectivity in the nucleus accumbens. Brain Struct Funct 104, 1 11 (2017).

116. Ungless, M. A. & Grace, A. A. Are you or aren’t you? Challenges associated with physiologically identifying dopamine neurons. Trends Neurosci 35, 422 430 (2012).

117. Pinault, D. A novel single-cell staining procedure performed in vivo under electrophysiological control: morpho-functional features of juxtacellularly labeled thalamic cells and other central neurons with biocytin or Neurobiotin. J Neurosci Meth 65, 113–136 (1996).

118. Morel, C. et al. Nicotinic receptors mediate stress-nicotine detrimental interplay via dopamine cells’ activity. Mol Psychiatr 36, 1418 (2017).

119. Manvich, D. F. et al. The DREADD agonist clozapine N-oxide (CNO) is reverse-metabolized to clozapine and produces clozapine-like interoceptive stimulus effects in rats and mice. Sci. Rep. 8, 3840 (2018).

